# Epigenetic regulations follow cell cycle progression during differentiation of human pluripotent stem cells

**DOI:** 10.1101/2020.06.26.173211

**Authors:** Pedro Madrigal, Siim Pauklin, Kim Jee Goh, Rodrigo Grandy, Anna Osnato, Daniel Ortmann, Stephanie Brown, Ludovic Vallier

## Abstract

Most mammalian stem cells undergo cellular division during their differentiation to produce daughter cells with a new cellular identity. However, the cascade of epigenetic events and molecular mechanisms occurring between successive cell divisions upon differentiation have not yet been described in detail due to technical limitations. Here, we address this question by taking advantage of the Fluorescent Ubiquitination-based Cell Cycle Indicator (FUCCI) reporter to develop a culture system allowing the differentiation of human Embryonic Stem Cells (hESCs) synchronised for their cell cycle. Using this approach, we have assessed the epigenome and transcriptome dynamics during the first two divisions leading to definitive endoderm. We first observed that transcription of key markers of differentiation occurs before division suggesting that differentiation is initiated during the progression of cell cycle. Furthermore, ATAC-seq shows a major decrease in chromatin accessibility after pluripotency exit indicating that the first event of differentiation is the inhibition of alternative cell fate. In addition, using digital genomic footprinting we identified novel cell cycle-specific transcription factors with regulatory potential in endoderm specification. Of particular interest, Activator protein 1 (AP-1) controlled p38/MAPK signalling seems to be necessary for blocking endoderm shifting cell fate toward mesoderm lineage. Finally, histone modifications analyses suggest a temporal order between different marks. We can also conclude that enhancers are dynamically and rapidly established / decommissioned between different cell cycle upon differentiation. Overall, these data not only reveal key the successive interplays between epigenetic modifications during differentiation but also provide a valuable resource to investigate novel mechanisms in germ layer specification.

## Introduction

Epigenome mapping during differentiation has been of great interest in recent years (Roadmap Epigenomics Consortium, et al, 2015). Indeed, histone modifications, enhancers’ formation and chromatin organisation have been directly linked to the establishment of cellular identity during differentiation of embryonic and somatic stem cells (González et al, 2015; Wang et al, 2015; Adam et al, 2015). However, these studies have often compared undifferentiated cells with cells produced several days after induction of differentiation thereby excluding early events inducing cell identity acquisition (Roadmap Epigenomics Consortium, et al, 2015; Tsankov et al, 2015; Dixon et al, 2015; Ziller et al, 2015). Furthermore, differentiation is often, if not systematically, associated with cell division in mammalian stem cells (Orford and Scadden, 2008). Consequently, mechanisms directing the acquisition of a new cellular identity are likely to be dynamically regulated during cell cycle progression. However, the study of these cell cycle related mechanisms is technically challenging *in vivo* due to ethical considerations especially in human, quantity of biological material and the complexity of cellular environment. In addition, stem cells grown *in vitro* are often heterogeneous in nature which renders difficult the analyses of cell cycle related mechanisms during differentiation. Together, these limitations might have concealed dynamic epigenetic regulations taking place during cell cycle progression upon differentiation. Here we use human Pluripotent Stem Cells (hPSCs) to address in part these questions. hPSCs can be grow *in vitro* almost indefinitely and maintain their capacity to differentiate into near homogenous population of primary germ layer progenitors using defined culture conditions devoid of unknown factor interfering with experimental outcome (Vallier et al 2009). Furthermore, previous studies already suggested transcription factors, epigenetic modifiers and signalling pathways that control cell fate choices in hPSCs (Yiangou et al, 2018). Accordingly, enhancer maps have been established very precisely during differentiation of hPSCs (Roadmap Epigenomics Consortium, et al, 2015; Wang et al., 2015) while the functions of transcription factors directing endoderm differentiation such as EOMES, GATA6, SOX17 and FOXA2 have been extensively studied (Teo et al 2011; Chia et al 2019; Fisher et al 2017; Séguin et al 2008; Genga et al 2019).

Finally, we and others have recently shown that human embryonic stem cells (hESCs) can be synchronised in different phases of their cell cycle using the FUCCI reporter system and then induced to differentiate into a near homogeneous population of endoderm cells (Singh et al 2013; Singh et al 2015; Pauklin and Vallier, 2013). Taking advantage of this approach, we further developed a culture system to differentiate hESCs synchronised for their cell cycle. This approach enabled us to analyse epigenetic modifications occurring during the two first cell cycles leading to endoderm, uncovering the early changes in the epigenome that direct the acquisition of a new cellular identity during differentiation.

## Results

### Culture system to differentiate cell cycle synchronised cells

We and others have shown that hESCs can only be induced to differentiate into endoderm during the G1 phase of their cell cycle (Pauklin and Vallier, 2013; Singh et al, 2013). Thus, we hypothesised that hESCs synchronised in G1 could differentiate homogenously while progressing through their cell cycle simultaneously. To confirm this possibility, a near homogenous population of hESCs was isolated in the early G1 phase (EG1-hPSCs) by cell sorting using the Fluorescent Ubiquitination-based Cell Cycle Indicator (FUCCI) reporter system (**Fig. 1a and Fig. S1a**) (Sakaue-Sawano et al., 2008; Pauklin and Vallier, 2013). FUCCI is a two-color (red and green) indicator that permits to follow cell cycle progression in live cells without the need chemical inhibitors (Pauklin and Vallier, 2013, Singh et al, 2016). The sorted cells were then replated in culture conditions inductive for endoderm differentiation previously validated with a diversity of hPSC lines (Touboul et al 2010; Cuomo et al, 2020). The resulting cells differentiated into a homogenous population of mesendoderm cells expressing the protein *T* after 36 hrs and definitive endoderm cells expressing the protein *SOX17* after 48hrs hrs (**Fig. 1b**). Further analyses revealed that EG1-hPSCs progressed through differentiation while being synchronised for their cell cycle for 24 hours which corresponds to the duration of the first cell cycle after induction of differentiation (**Fig. S1**) (Calder et al, 2013). Indeed, EG1-hPSCs progressed through S phase after 12 hrs and undergo a first division after 24 hrs to re-enter S phase 36 hrs after the induction of differentiation. The resulting mesendoderm cells undergo a second division around 48hrs to become endoderm cells which remain blocked in G1 until the end of our experimental time frame (72 hrs) (**Fig. 1a and Fig. S1a**). Considered together, these results confirm that our culture system can be used to differentiate cell cycle synchronised hESCs into endoderm cells, thereby providing a platform for investigating molecular regulations occurring during cell cycle progression upon differentiation.

**Figure 1:**
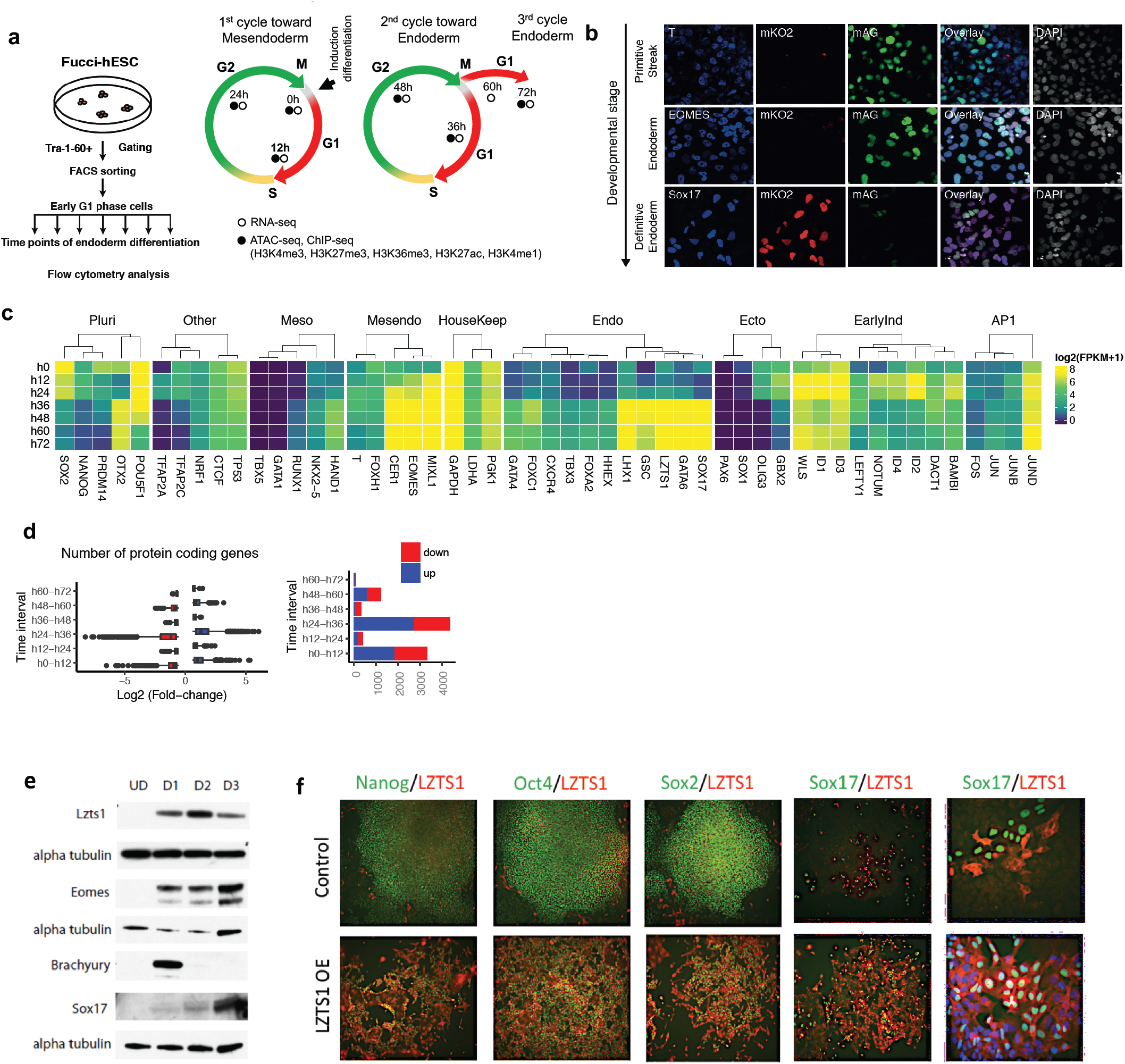
Cell cycle synchronization during differentiation of hESCs reveals that each cell division result into a new cellular identity. (**a**) Schematic representation of experimental setup to differentiate synchronised EG1-FUCCI hPSCs into endoderm and samples collected for sequencing analyses. Genome-wide data collected at each time point is indicated. (**b**) Immunofluorescence analyses showing expression of primitive streak (*T*, 36 hrs), early-endoderm (*EOMES*, 48 hrs) and definitive endoderm (*SOX17*, 72hrs) markers in FUCCI-hESCs differentiating into endoderm after synchronisation. (**c**) Gene expression profile analyses showing selected genes that are differentially expressed during cell cycle progression upon differentiation. (**d**) Number of differentially expressed genes (FC≥1.5; FDR≤0.01) (right) and distribution of log_2_ fold-changes (left) during cell cycle progression upon differentiation. (**e**) Western blot analyses showing the expression of mesendoderm markers (T), endoderm markers (SOX17, EOMES) and LZTS1 during differentiation of hPSCs into endoderm. (**f**) Immunostaining showing the expression of pluripotency markers (NANOG, OCT4, SOX2), endoderm markers (SOX17) and LZTS1 in hPSCs (control) and hPSCs overexpressing LZTS1 (LZTS1 OE).

### Gene expression marking differentiation starts before cell division

Taking advantage of our culture system, we decided to investigate the dynamic of epigenetic changes occurring during progression of cells cycle upon differentiation. For that, we performed genome-wide analyses including RNA-Seq, ATAC-Seq, histone marks ChIP-Seq (H3K4me3, H3K27me3, H3K27ac, H3K4me1, H3K36me3) on EG1-hPSCs differentiating for 12hrs (Early/Late G1 of 1st cell cycle); 24 hr (S/G2/M of 1st cell cycle); 36 hrs (S/G2/M phase of 2nd cell cycle), 48 hrs (end of second cell cycle) and 60/72 hrs (G1 of 3rd cell cycle) (**Fig. 1a**). Quality and reproducibility of datasets were confirmed, demonstrating the robustness and reproducibility of our approach (**Methods**). We first decided to characterise gene expression in hPSCs differentiating into endoderm. In agreement with previous reports, RNA-Seq analyses revealed a precise timing and succession of gene expression (**Fig. S1b and S1c**) upon endoderm differentiation (Chu et al 2016; Cuomo et al 2020; Pauklin et al, 2016). Of particular interest, *SOX2* represent the first pluripotency marker to decrease upon induction of differentiation in agreement with its known function in repressing mesendoderm specification (Wang et al, 2012). Decrease in *SOX2* was rapidly followed by the induction of mesendoderm markers at 12 hrs starting with *T*, and followed by *MIXL1* and *EOMES* (**Fig. 1c**). Interestingly, mesendoderm markers (*T, EOMES, MIXL1*) start to increase before 24hrs and thus before the first division. On the other hand, pluripotency markers (*OCT4/POU5F1, NANOG*) continue to be expressed after the first cell division (24hrs) thereby confirming that pluripotency factors have a role in inducing mesendoderm specification (Radzisheuskaya et al 2013). However, pluripotency markers strongly decreased during the second cell cycle (36 hrs) while endoderm markers (*GATA4/6* then *SOX17* and *FOXA2*) were induced, suggesting that endoderm specification could start before the second division. These observations confirm that the first cellular division upon differentiation produces mesendoderm cells (*OCT4*+/*T*+/*SOX2*-), while the second division gives rise to endoderm cells (*SOX17*+/*GATA6*+/ *OCT4*-). Interestingly, gene expression mostly varies during G1 phase transition (0-12hrs or 24-36hrs and 48-60hrs) with low transcript change (both in number of genes and the fold-change) during S phase (**Fig. 1d**). Thus, each specification phase (pluripotency to mesendoderm, and mesendoderm to endoderm) starts before the cellular division which ultimately produces a new cell type.

### Cell cycle synchronisation of differentiating cells reveals new markers for endoderm differentiation

To further investigate our RNA-Seq data and identify new markers for endoderm differentiation, we performed *k*-means clustering of 6317 differentially expressed genes (0h, 12h, 24h, 36h, 48h, 60h and 72h). This analysis revealed 13 gene clusters whom expression dynamically changes with differentiation, with several gene clusters displaying dynamic and transitory expression (**Fig. S1d and S1e**). *NOTUM, WLS, BAMBI, CRABP1/2, DACT1, NKD and ID1/2/3/4* were immediately and transitorily expressed upon induction of differentiation (clusters 5, 7; **Fig. S1e and Table S1**). Except for *ID1*, the expression of these genes have not been described previously in the context of endoderm differentiation (Chu et al, 2016). Their rapid decrease after induction might have masked their expression in previous studies while suggesting that they could be involved specifically in the earliest step of differentiation. We also observed gene clusters with expression patterns similar to those of known master regulators of endoderm specification such as SOX17, and thus hypothesised that such genes could have an important role. LZTS1 appeared to be particularly interesting since it is expressed in the mouse primitive streak, it is bound by EOMES (Teo et al 2011), and its expression is strongly induced after 36hrs of differentiation, following closely GATA4/6 and SOX17 induction (Cluster 3; **Fig. S1e and Table S1**). Rapid upregulation of LZTS1 after 36hrs of differentiation was confirmed by western blot (**Fig. 1e**) while gain of function experiments in hPSCs showed that *LZTS1* overexpression increased expression of *SOX17* thereby suggesting a function in endoderm specification (**Fig. 1f**). Taken together, these analyses reveal distinctive transcriptional waves of transcription during early cell fate decisions which include previously unknown potential regulators of endoderm differentiation.

### Early changes in gene expression are not associated with modification of chromatin accessibility

Based on the dynamic expression pattern observed in our transcriptomic analyses, we investigated whether the observed gene expression changes were associated with chromatin modifications. For that, we performed ATAC-Seq (Buenrostro et al., 2013) analyses at different time points during cell cycle progression upon differentiation (**Fig. 1a**). Computational analyses revealed chromatin accessibility changes in 31,018 genomic regions, out of 253,349 analysed, between consecutive intervals of the time-series (|*FC*|≥2; adj-*P*≤10^−4^). These changes mainly occur in regions containing protein coding-genes and long intergenic noncoding RNAs (**Fig. S2a**). Generally, chromatin accessibility changes in intergenic, intronic or regions located upstream of genes occur independently of the cell cycle phase or stage of differentiation (**Fig. S2b**). Indistinctly of genomic annotation, chromatin mostly compacts during the first and last G1 phase, while changes between 24-48h were dominated by an increase in accessibility (**Fig. 2a**) suggesting that different epigenetic regulations could occur between the two cell cycles leading to endoderm. Despite the increase of chromatin accessibility at intermediate stages, average chromatin accessibility was drastically reduced over the three days (**Fig. 2b**) confirming that hESCs display an open chromatin landscape which could provide the necessary opportunity for differentiation induction (Dalton, 2015). Of note, genes close (up to 10kb) to accessibility-decreased regions in the first G1 phase were enriched in GO terms of ‘nervous system development’ (*P* = 2.5e^-18^) and ‘Axon guidance’ (*P* = 0.00036) suggesting that restriction of cell fate decision toward the neuroectoderm pathway through chromatin reorganisation could be the first event of endoderm differentiation.

**Figure 2:**
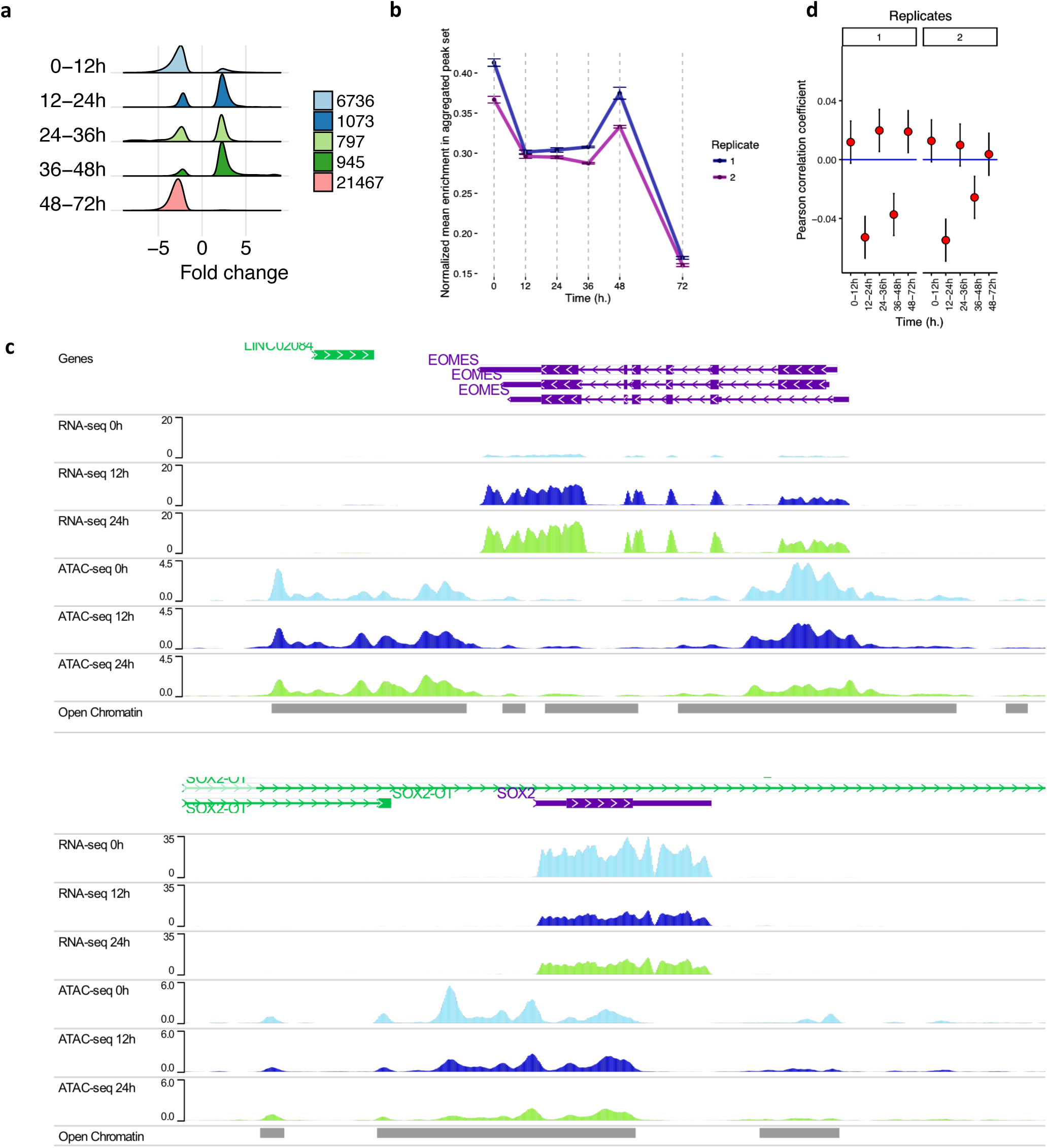
Chromatin accessibility during differentiation of hPSCs synchronised for their cell cycle. (**a**) Number of regions showing significant chromatin accessibility increase (“opening” ; FC>2 and adj-*P*≤10^−4^) or decrease (“closing” ; FC<2 and adj-*P*≤10^−4^) during progression of cell cycle during differentiation of cell synchronised EG1-FUCCI hPSCs (0-12hrs: G1 to S during 1^st^ cell cycle; 12-24hrs: S to G2/M during 1^st^ cell cycle; 24-36 hrs: 1^st^ cell division to S during 2^nd^ cell cycle; 36-48hrs: S to G2/M during 2^nd^ cell cycle; 48-72 hrs: 2^nd^ cell division to G1 of 3^rd^ cell cycle). (**b**) Normalised mean read-enrichment in ATAC-seq consensus peaks during endoderm differentiation of EG1 FUCCI-hPSCs. (**c**) ATAC-Seq and RNA-Seq genome browser track visualisation showing change in EOMES and SOX2 (GENCODE v29) expression and associated chromatin status during the first 36h of differentiation. Data tracks shown correspond to the first replicates. Open chromatin represents a merged consensus during differentiation. (**d**) Pearson’s product-moment correlation between log_2_ Fold-change in RNA (reported by DESeq2) and fold-change of the maximum normalized ATAC-Seq signal in the two replicates in a region 10kb upstream of the promoter of protein-coding genes. 95 percent confidence interval is shown.

Next, we correlated changes in gene expression with variant chromatin accessibility. We found that while increase in *MIXL1, T* and *EOMES* expression took place in invariant chromatin regions during the first 12h, *SOX2* chromatin accessibility decreased in concert with its expression (**Fig. 2c**). Similarly, change in transcription at 12-24h for *MIXL1, GATA6, GSC, CER1, T, LZTS1*, and *SOX17* did not display modification of chromatin accessibility (**see: http://ngs.sanger.ac.uk/production/endoderm**). Correlation between fold change in ATAC-Seq and RNA-Seq was only significant, and negative, in time intervals 12-24h, 36-48h which are associated to S/G2 (*P*<0.005, Pearson’s product-moment correlation; **Fig. 2d**) which presents little transcriptional change (**Fig. 1d**). Similar analyses on protein coding genes confirm the lack of correlation between change in chromatin and transcription (**Fig. S2c**) thereby suggesting a weak coupling between transcriptional induction and chromatin reorganisation during early differentiation. Taken together, these results suggest that chromatin organisation changes rapidly during differentiation to block alternative cell fate during cell cycle progression but not to enable the expression of genes marking mesendoderm.

### Footprinting analyses show dynamic regulation of transcription factor binding during cell cycle progression upon differentiation

To further the understanding of the dynamic regulation in this system, we identified overrepresented transcription factor (TF) DNA binding motifs in dynamic chromatin accessibility regions observed during differentiation. Motif enrichment analyses revealed that regions with increased accessibility contains motifs for effector of signalling such as ACTIVIN/NODAL (SMAD2), BMP (SMAD1) and WNT (TCF4/12) which are known to drive endoderm differentiation (**Fig. S3a**). On the other hand, CTCF binding motif was highly enriched in regions with decreasing chromatin accessibility during G1 phase (0-12h, 48-72h, **Fig. S3a**). Similarly, Activator protein 1 (AP-1) motifs (JUND, JUN, FOS, JUNB) were the most significantly enriched in regions displaying a decreased ATAC-Seq peaks associated with chromatin compaction (24-36h and 36-48h, **Fig. S3a**). Nonetheless, some TFs seem to have opposite function at different phases of cell cycle progression during differentiation. As an example, CTCF binding motifs were also strongly associated with regions displaying an increased accessibility in S phase progression (36-48h, **Fig. S3a**). Taken together, these motif enrichment analyses suggest that different TFs could have specific functions during specific phase of the cell cycle during the progression of differentiation.

To further identify active DNA regulatory elements, we performed digital genomic footprinting as a proxy for transcription factor binding. This approach locates depleted narrow regions within open chromatin created by TFs binding to DNA and preventing Tn5 cleavage. Genomic footprinting was first introduced in DNase-seq, and it has been performed with success in ATAC-seq (Corces et al, 2018). Using this approach, we identified more than 2 million reproducible, non-redundant, bias-corrected, putative TF binding sites, detected by two independent software tools, and associated to genes that at the same stage were expressed at least 1 FPKM (**Methods**; **Fig. S3b**). 75% of our 0h/EG1 footprints co-localize with H7-hESC ENCODE footprints (Neph et al., 2012) confirming the accuracy of our analyses (**Methods**). Interestingly, these analyses reveal that unique regions of TF footprints were annotated more to promoter than intronic regions in hESC (0h) and DE (72h), but vice versa in intermediate time-points (**Fig S3c**). This suggests that “stable” cell identities have a transcriptional network relying mostly on promoters, whereas in cell transitions rely more on enhancer rewiring.

We next decided to identify transcription factors for which the binding results in major changes in chromatin organisation in subsequent phase of the cell cycle. For that, we implemented a predictive model to systematically study TF action in chromatin accessibility dynamics. A functional linear model with a scalar response was used to predict the *log*_*2*_|*FC*| of ATAC-seq chromatin accessibility in the peaks between two consecutive stages with |*FC*|*>*1.5 for TF footprints that overlap those regions (**Methods**). The linear model was only considered when at least 15 footprints overlapped the regions of interest. The analysis was performed using as a reference footprints detected either at initial or final time points. We assessed the quality of the fit for each TF by computing the multiple square correlation (*RSQ*) and *F*-ratio (Wald test). *P*-values for the *F*-statistic were calculated to evaluate whether the fit to the data is better than what we would expect by chance. We observed low *RSQ* suggesting that DNA-binding of each TF in isolation does not predict well ATAC-seq change confirming that combinatorial binding of several factors is necessary in the context of chromatin (Reiter et al, 2017). Nonetheless, we obtained statistically significant results for footprints of diverse TFs including for example at 36h FOXA2, SMARCC1, CTCF, TP53 which were the best predictors for the decrease in chromatin accessibility occurring at 48h (for detailed analyses see supplementary results). Footprints for proteins of the AP-1 complex found at 48h were top predictors of chromatin compaction between 24-48h (**Fig. 3b**), in agreement with our DNA motif enrichment analyses (**Fig. S3a**). To investigate these observations further and reveal TF dynamics in a statistically robust manner, differential ATAC-seq footprinting was performed with Wellington_bootstrap (**Fig. 3a**), which is able to account for different sequencing depths of the datasets (Piper et al., 2015). Most significant overrepresented TFs were NRF1 (12h), TP53 and TFAP2A/C (24h), and JUND (36h), a member of the AP-1 transcription factor complex (**Fig. 3c**). Reanalysis of the data with a recent algorithm for bivariate genomic footprinting showed similar results and uncovered the GATA motif in increased chromatin accessibility at 48h (**Fig. 3d**). These results confirm that different combinations of transcription factors could organise chromatin at specific phases of the cell cycle upon differentiation. They also reveal significant effects of specific TF binding in chromatin modulation and suggest that AP-1 factors could have a key function in modulating chromatin accessibility after the first cell division.

**Figure 3:**
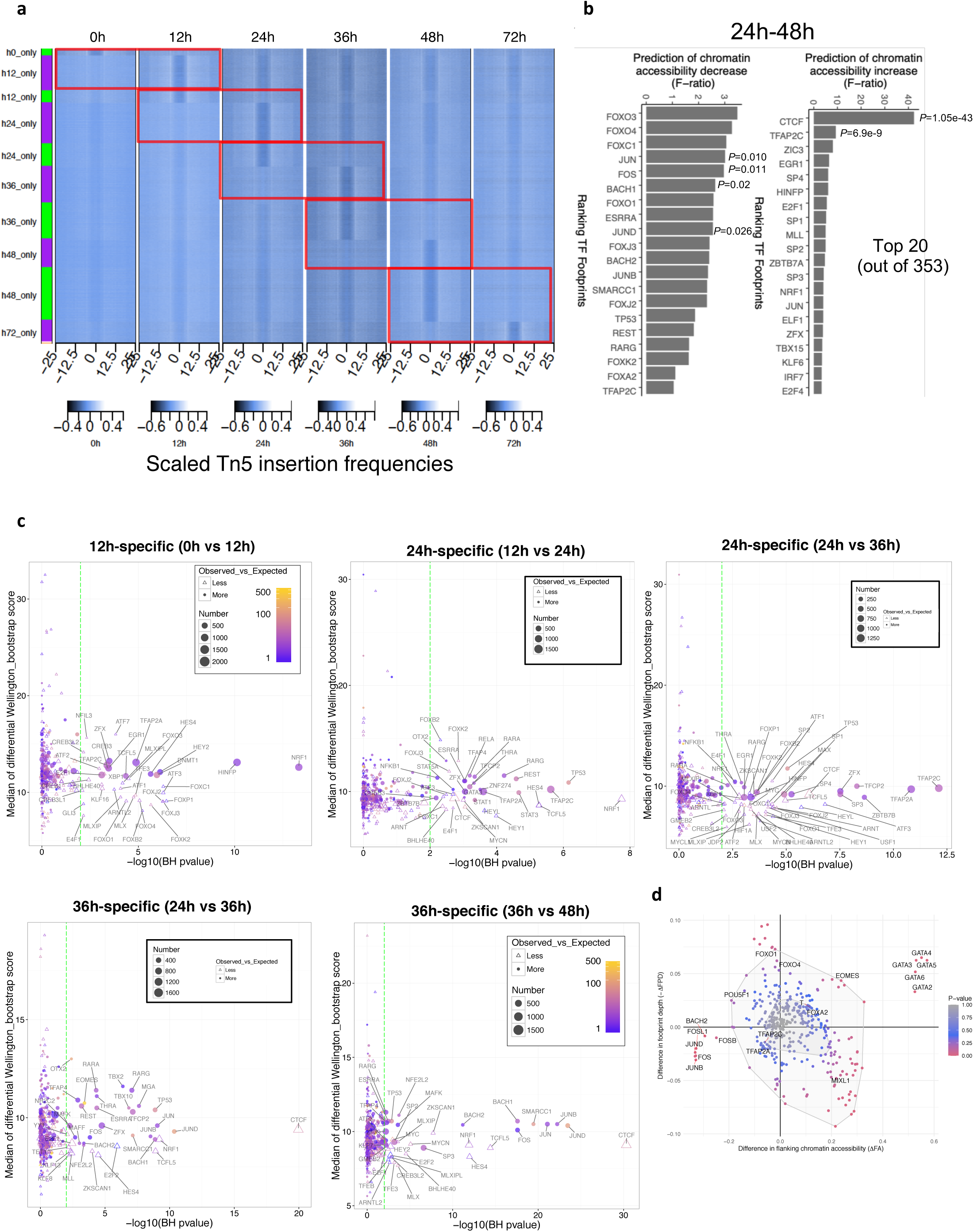
Digital genomic footprinting reveals the dynamic activity of key transcription factors during progression of cell cycle upon differentiation. (**a**) Heatmap of normalized Tn5 insertions for differential footprints (Wellington bootstrap score *S*>20). Red rectangles drawn indicate the results for each pairwise comparison. (**b**) Top 20 best predictors of a functional linear model using TF footprints at 48h to model chromatin accessibility change between 24-48h. *P*-value associated with the *F*-ratio, to evaluate whether the fit to the data is better than what we would expect by chance, is shown for selected TFs. (**c**) Overrepresentation analysis for TFs associated to differential footprints shown in (b). Each differential footprint was first matched to the consensus list of footprints detected by FootprintMixture and Wellington (see Methods). *P*-values (Chi-squared test; Benjamini–Hochberg multiple hypothesis testing correction) indicate if proportions of footprints at each time point for a TF are significantly different when comparing differential footprints and the total number of footprints at this time point. Only TFs with adjusted *P*≤0.01 have been labelled. (**d**) Bag plot depicting changes in flanking chromatin accessibility (ΔFA) and footprint depth (ΔFPD) in ATAC-seq of endoderm differentiation between 24hrs and 48hrs in human motifs. Statistically significant change in FA/FPD was evaluated by chi-square distribution. Motifs of genes that were not differentially expressed during endoderm differentiation (RNA-seq) were removed. Outlier ARID5A (ΔFA=0.15, ΔFPD=0.27) was also removed from the plot.

### MEK/ERK/p38/AP-1 signalling pathways control different steps of endoderm differentiation

To validate the functional relevance of computational ATAC-seq analyses, we investigated the importance of AP-1 signalling pathways in DE differentiation. For that, hPSCs were differentiated in the presence of inhibitors for three different pathways which are known to control the AP-1 complex: MEK1/2 (U0126-EtOH), p38-MAPK (SB203580) and JNK (JNK-IN-8) (**Fig. 4a**). These analyses revealed that p38-MAPK inhibition blocked the expression of DE markers (*GATA4, SOX17, CDX2* and *FOXA2*, **Fig. 4b-d and fig. S3d,e**) without affecting the increase of primitive streak markers (*T, EOMES*, **Fig. 4b and Fig. S3d**) or the decrease of pluripotency markers (*OCT4*/*POU5F1, NANOG, SOX2*, **Fig. 4b and Fig. S3d**) while promoting mesoderm markers expression (*MIXL1, NKX2*.*5*, **Fig. 4b and Fig. S3d**). On the other hand, JNK inhibition marginally increased expression of endoderm markers and inhibition of ERK5/MAPK blocked DE differentiation while limiting pluripotency markers decrease (**Fig. 4b-d and fig. S3d, e**). Thus, these data suggest that the p38-MAPK-AP1 pathway is necessary for endoderm specification and for full dismantling of the pluripotency network. Combined together, these experiments and the computational analyses shed light on the different effects exhorted by the signalling controlling the AP-1 complex. MAPK-ERK seems necessary to exit pluripotency during endoderm specification (Singh et al., 2012; Na et al., 2010) while MAPK-p38 promotes endoderm induction by inhibiting mesoderm. This diversity of functions could be achieved by dynamic interactions between the members of the AP-1 at different stages of cell cycle progression upon differentiation.

**Figure 4:**
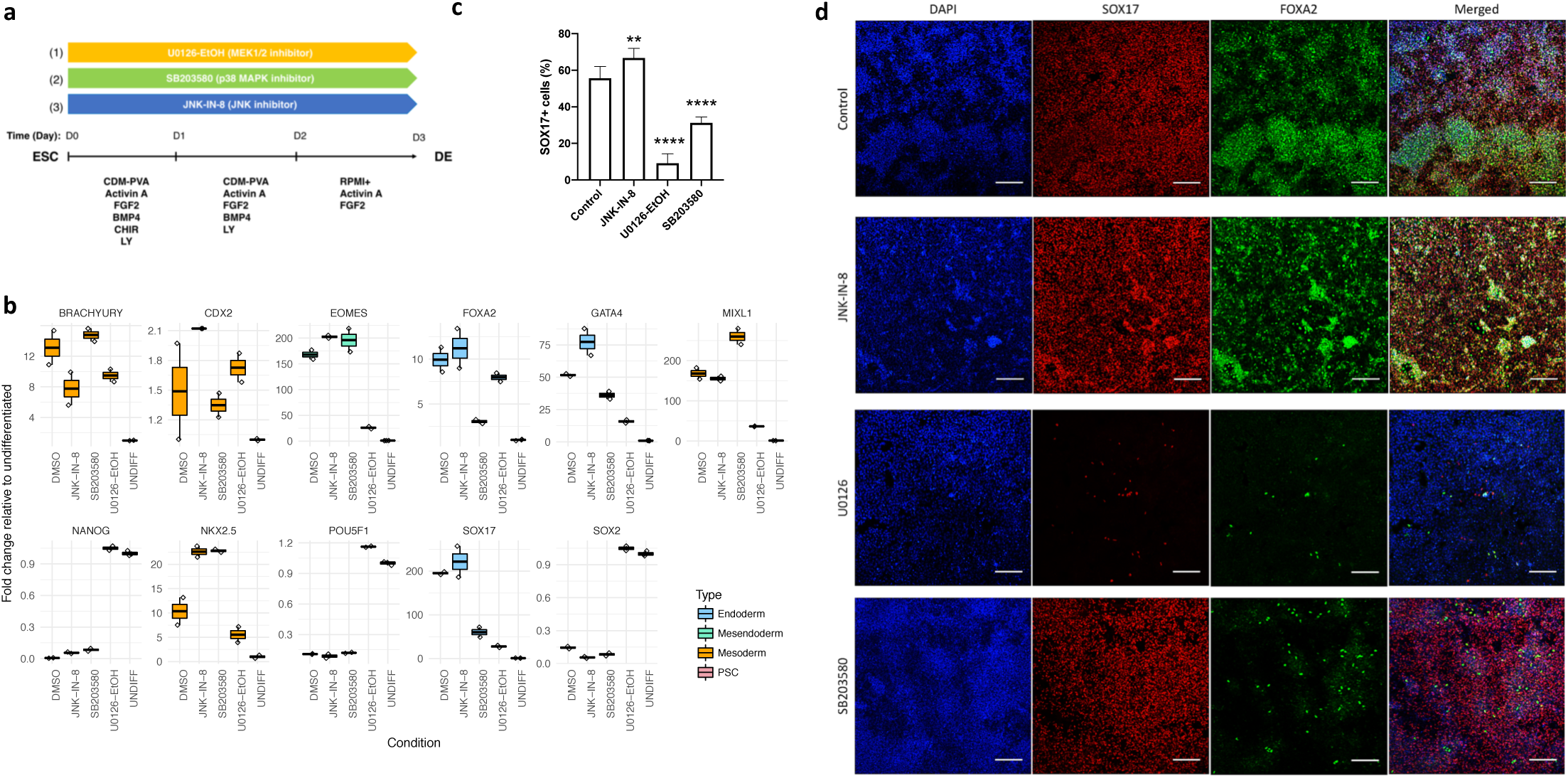
Inhibition of AP-1 complex blocks endoderm differentiation of human pluripotent stem cells. (a) Schematic representation of the experimental plan to characterise the functional relevance of MEK1/2, JNK and p38 pathways in endoderm differentiation. hPSCs were grown for 3 days in culture conditions inducing endoderm differentiation in the presence of small-molecule inhibitors. (**b**) Q-PCR, (**c**) FACs and (**d**) immunostaining were performed after 3 days for pluripotency markers (POU5F1/OCT4, NANOG, SOX2), mesendoderm/mesoderm markers (BRACHYURY/T, NKX2.5, CDX2, MIXL1) and endoderm markers (SOX17, EOMES, GATA4, FOXA2). For QPCR, the average of 2 separate experiments including 6 different biological replicates is provided with STD. For FACs, one-way ANOVA was performed followed by Dunnett’s multiple comparisons test where each of the 3 treatment conditions were compared against the control. For immunostaining, ccale Bar 200µm.

### Cell cycle progression upon differentiation is associated with dynamic histone modifications

To further investigate the dynamic change in epigenetic state occurring during differentiation, we performed ChIP-seq analyses for five histone modifications (H3K4me3 for active promoter region, H3K4me1 and H3K27Ac for enhancer, H3K27me3 for repressed region, and H3K36me3 for gene bodies of actively transcribed genes). We identified differential ChIP-seq regions between consecutive stages, and observed that transition from pluripotency to DE involved several waves of histone modifications (**Fig. 5a**). H3K4me3 and H3K27ac marks were the most dynamic during differentiation. H3K4me3 decreases during the start of differentiation (0-12h) in agreement with our ATAC-seq analyses suggesting that the first step of differentiation consist in inhibiting alternative fate choice. In addition, early developmental loci rapidly acquire H3K4me3 and H3K27ac marks during S phase (12-24h) confirming that epigenetic modifications characterising differentiation could start before the first division. Accordingly, regions surrounding key genes such as *T, GSC, SOX17, EOMES, MIXL1, GATA6, GATA4, LHX1, FOXA2* acquire H3K27ac rapidly upon differentiation (0-12hrs, **Fig. 5b**; see http://ngs.sanger.ac.uk/production/endoderm). Major acquisition/increase of enhancer marks (H3K27ac and H3K4me1) also occurs before the second division (24-36h) or before the third cell division (48-72h) (**Fig 5a**). Accordingly, clustering of Jaccard indexes using all the reproducible peaks in ChIP-seq and ATAC-seq (those with >100 regions) to quantify the overlap of chromatin marks during differentiation confirmed that enhancer marks (H3K27ac, H3K4me1) and open chromatin are very dynamic (**Fig. 5c**). In addition, despite the lack of quantitative peak changes, we could observe that H3K27me3 chromatin regions at 0h and 72h were most dissimilar, and that H3K36me3 peak sets clustered into two groups: before and after the first cell division (**Fig. 5c**). Change in H3K27ac/H3K4me3 and H3K27ac/H3K4me1 seem to associated more than any other marks, while change in chromatin accessibility was also linked with the deposition of these marks. As expected H3K27me3 and H3K36me3 or H3K27me3 and H3K27ac were mutually exclusive.

**Figure 5:**
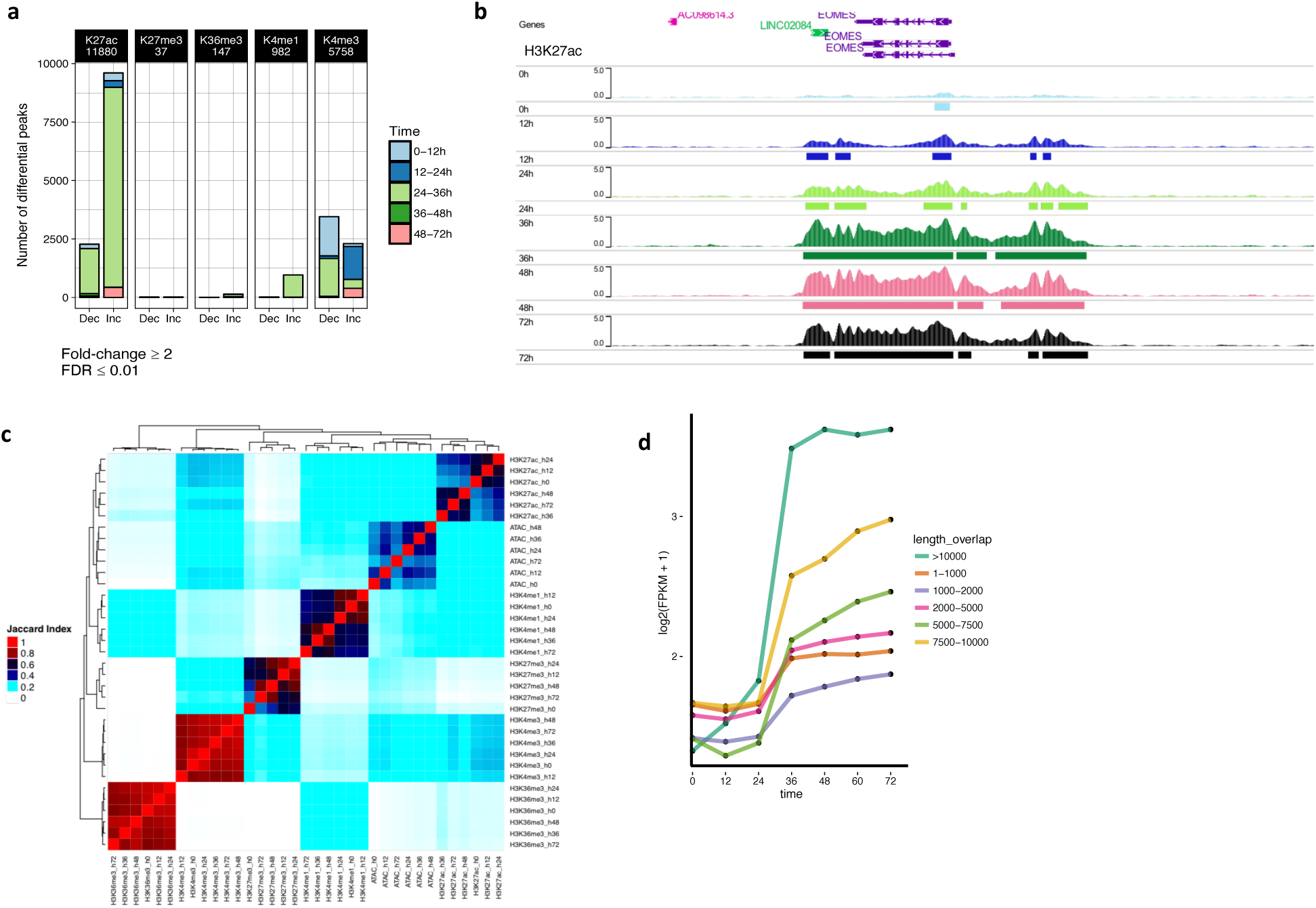
Chromatin modifications occurs dynamically during cell cycle progression upon differentiation. **(a)** Number of genomic regions containing dynamic chromatin marks in consecutive time points based on histone ChIP-seq. **(b)** H3K27ac ChIP-seq for *EOMES* locus (GENCODE v29; first replicate shown). **(c)** Hierarchical clustering of Jaccard Index values obtained for overlaps between ChIP-seq and ATAC-seq regions. **(d)** Gene expression versus length of the direct overlap between H3K4me1 and H3K27ac both increasing 24-36h (protein coding genes 10kb around).

We then decided to correlate the change in gene expression with histone mark acquisition focusing on H3K4me1 and H3K27ac since 784 out of 954 H3K4me1 increased regions were coincident with H3K27ac increase in 24-36h. We observed that, when both marks increased simultaneously, genes in close proximity increased its expression proportionally to the size of the overlap between the two chromatin marks (**Fig. 5d**). Interestingly, this correlation starts immediately upon differentiation suggesting that tissue super-enhancer (SE, marked by H3K27ac and H3K4me1 spanning an intersection of >3 kb) (Parker et al 2013; Hnisz et al 2013) seems to be initiated before division during the first S phase to fully established only after the first division thereby following the induction of expression for key developmental regulators (**Fig. 1c**). Taken together, these data suggest the existence of hierarchy between histone marks during differentiation. H3K27ac appears on key genes marking nascent mesendoderm before or during the induction of their expression immediately after the G1 phase. On the other hand, H3K27me3 and H3K36me3 are less dynamically regulated and seem to change after gene expression and/or chromatin organisation.

### Transient super-enhancers after each cell division define cell identity

Next, we decided to use our ChIP-Seq data to further characterise the enhancer regulatory landscape defining cellular identity acquisition. For that, we identified at each time point poised enhancers (PEs, H3K27ac-, H3K4me1+), active enhancers (AEs, H3K27ac+, H3K4me1+ < 3kb), and Super Enhancers (SEs, H3K27ac+, H3K4me1+ > 3 kb). As H3K4me3 has been detected in promoters (Atlasi and Stunnerberg, 2017), we subdivided AEs and SEs as proximal or distal according to presence of H3K4me3 (**Fig. 6a**). Distal AEs and Distal SEs were the most dynamic regulatory regions with their number increasing immediately after differentiation to peak at 36hr (**Fig. 6b**) and then decreasing until 72hrs. Proximal AEs and poised enhancers were relatively stable during cell cycle progression. These results confirm that SE establishment starts immediately upon differentiation to be consolidated after the first division to establish a new cellular identity. We then perform clustering of Jaccard indexes to identify similarities between regulatory regions during progression of cell cycle. Using this approach, we observed that poised enhancers partially converted to distal AEs after one division, especially at 36h. More interestingly, distal SEs at 0h and 72h were less similar to those at 12/24h and 36/48h, respectively suggesting that acquisition and partial loss of SEs is occurring actively at each cell division resulting in a new cellular identity (**Fig 6c**). Nonetheless, while SEs were different before and after first cell division, SEs at 48hrs were a subset of those established at 36h (**Fig. 6d**). For example, key mesendoderm/endoderm loci such as *CER1, EOMES, LZTS1, MIXL1* contain several SEs established and maintained after first cell division (*FC*>4; *P*≅0) (http://ngs.sanger.ac.uk/production/endoderm). Other SEs at 36h include *EOMES, LHX1, GSC, ZIC3, OTX2, DKK1, WLS, MYC, HAND1, WNT3*, as well as some members of the MAPK family, among others. Thus, SEs seem to be established following two different regulatory mechanisms between the first and second division (**Fig. 6e**). A large number of new SEs seem to be established after the first division at the mesendoderm stage, while SEs characterising endoderm cells are simply maintained during the second division and mesoderm specific enhancers are progressively lost. Thus, cellular identity acquisition is achieved by the creation or selection of super enhancers starting before division thereby suggesting the existence of different epigenetic regulations between successive cell cycles upon differentiation.

**Figure 6:**
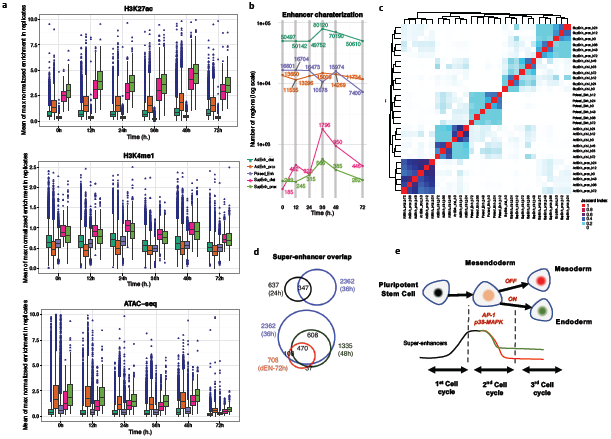
Epigenetic dynamics during cell cycle progression upon differentiation reveals super-enhancers assembly and loss to establish a new cellular identity. (a) Distribution of H3K27ac, H3K4me1, and ATAC-seq signal in the enhancer regions shown in (b). (b) Enhancer classification during endoderm differentiation: ActEnh_dist (H3K27ac+, H3K4me1+, H3K4me3-, < 3 kb); ActEnh_prox (H3K27ac+, H3K4me1+, H3K4me3+, < 3 kb); Poised_Enh (H3K27ac-, H3K4me1+); SupEnh_dist (H3K27ac+, H3K4me1+, H3K4me3-, > 3 kb); SupEnh_prox (H3K27ac+, H3K4me1+, H3K4me3+, > 3 kb). (c) Hierarchical clustering of Jaccard Index values obtained for overlaps between different enhancer regions. (d) Endoderm specific super-enhancers are a subset of those established at 36h. (e) Schematic model of super-enhancer establishment during endoderm differentiation at each consecutive cell division.

## Discussion

Our data provide a comprehensive epigenetic resource outlining the progressive acquisition of endoderm identity upon consecutive cell divisions. Our analyses also show that early differentiation follows gradual and stepwise transitions following cell cycle. Indeed, differentiation is initiated in the G1 phase and expression of differentiation of mesendoderm markers is initiated before the end of S phase. This timing implies that the first step of mesendoderm specification could be achieved through the re-organisation of the pluripotency network induced by changing culture conditions. Nonetheless, we did observe the transitory induction of early genes upon induction of differentiation. Most of these genes are known to control signalling pathways such as WNT for *NOTUM* or BMP for *BAMBI* (De Robertis and Kuroda, 2004; Zhang et al, 2015; Malaguti et al, 2019). In addition, some of these genes (*ID1*/*3*/*4*) are controlled by BMP signalling which is added in our culture condition. Thus, these genes might only mark a response to the change in culture conditions rather than actively direct differentiation. Alternatively, they could also repress neuroectoderm differentiation. Accordingly, we found that initiation of differentiation is associated with a decrease in chromatin accessibility around neuronal related genes, suggesting that one of the first event of differentiation is to block alternative cell fate specifications. Moreover, the rapid decrease in *SOX2* expression is likely to play a key function in this process. Indeed, *SOX2* is a key regulator of neuroectoderm specification while also being a key inhibitor of mesendoderm. On the other hand, we could not detect major changes in chromatin organisation associated with induction of mesendoderm markers suggesting that the corresponding genomic regions are already accessible in pluripotent stem cells. This invariant chromatin accessibility could imply that the reorganisation of the pluripotency network merely needs to activate the transcription of mesendoderm genes without major changes in chromatin accessibility. This mechanism will enable fast and rapid induction of differentiation following change of culture conditions. Such process is likely to be essential in a constantly changing environment such as the gastrulating embryo.

Interestingly, tissue specific super-enhancers start to be established before division to be fully established after division. Thus, transcription factors for which expression is induced in S phase become fully active in the following G1 phase. Alternatively, some of these factors could also act as mitotic bookmarking factors to program the identity of the newly create daughter cells before cellular division (Festuccia et al, 2017). Similar mechanisms seem to exist between the first division producing mesendoderm cells and the second division producing endoderm cells. However, the second division is mainly associated with a reduction in the number of super-enhancers. Thus, differentiation could start with the establishment of diversity of tissue-specific super enhancers which are then successively selected to enable production of more committed cell (either endoderm or mesoderm). Further functional studies will help to identify the mechanism by which mesendoderm and endoderm transcriptional networks could achieve divergent activities.

The JNK–JUN pathway inhibits exit from the pluripotent state via JUN binding on pluripotency enhancers with OCT4, NANOG, SMAD2 and SMAD3 (Li et al, 2019b). By digital genomic footprinting time-, cell phase-specific cis-regulatory elements related to the MAPK signalling pathway were identified to participate in the closure of the pluripotency network. Our results suggest that the AP-1 complex is likely to play a key part in the establishment and resolution of the mesendodermal network toward endoderm. We found AP-1 complex-associated TFs as best predictors of chromatin accessibility change in 24-48h and proved that p38/MAPK is indispensable for endoderm specification by using small molecule inhibitors. Thus, control of AP-1 activity by p38/MAPK which is likely activated by addition of FGF and PI3Kinase inhibition in our culture system is essential for the first initial differentiation. Transient AP-1-bound enhancers have been found during Oct4/Sox2/Klf4/cMyc (OSKM)-mediated cell reprogramming, linked to the extinction of the somatic transcriptional network (Madrigal and Alasoo, 2018). Thus, AP-1 could have a broad function beyond early human development to eliminate chromatin roadblocks and facilitate cell-state transitions.

Importantly, other transcription factors expressed in hPSCs such as NRF1, TFAP2C, TP53 (Wang et al, 2017) and CTCF seem to have a key function in the early induction of mesendoderm specification. CTCF is not only pivotal for the orchestration of topologically associated domains and loops, but also it displays cell-cycle dependent DNA-binding (Oomen et al, 2019). NRF1 is a methylation-sensitive TF that could behave as a pioneer factor during early differentiation (Mayran and Drouin, 2018), while TFAP2C could act as a settler TF (Slattery et al, 2014) binding all accessible sites before the first cell division. Moreover, TFAP2C has been recently involved chromatin accessibility dynamics in early ectoderm differentiation from hPSCs (Li et al., 2019). Thus, the mechanisms uncovered by our study could also apply to alternative germ layers. Nonetheless, further investigation will be necessary to understand the functions and the interplays of these different factors in mesendoderm specification and during each cell cycle leading to endoderm cells.

Finally, our results also underline the importance to include cell cycle progression in epigenetic and transcritpomic analyses. Indeed, our ChIP-seq profiling showed that during transition of cell-identity H3K27ac presents its highest increase at 36h resulting in the apparition of novel super-enhancers after the first cell division. However, most of them are lost at Day 3 and thus could not be detected in previous studies using unsynchronized cells (Tsankov et al., 2015). Furthermore, adding a cell cycle dimension in our analyses allowed to order molecular event which successively lead to change in cellular identity. Of note, some of the mechanisms uncovered are likely to be conserved *in vivo*. As an example, the formation of super-enhancers and fate restriction by chromatin closure are in agreement with recent studies in mouse post-implantation embryos (Argelaguet al., 2019). Thus, these regulatory mechanisms shaping the epigenome are likely to be relevant not only for developmental processes but also for other stem cells involved in normal homeostasis and diseases.

## Acknowledgements

We are grateful to Dr Irina Mohorianu, Dr Rute Tomaz and Marion Perrin for helpful comments, as well as to Sapna Vyas, Nathalie Smerdon and Siobhan Austin-Guest (Wellcome Sanger Institute) for help with sample management. This work was supported by the European Research Council Grant Relieve-IMDs and New-Chol (L.V., P.M, R.G.), the Cambridge Hospitals National Institute for Health Research Biomedical Research Center (L.V.) and a core support grant from the Wellcome Trust and Medical Research Council to the Wellcome Trust – Medical Research Council Cambridge Stem Cell Institute. S.P. was funded by a Federation of European Biochemical Societies long-term fellowship and a InnovaLiv EuFP7 grant. We thank the staff in the Sequencing core facilities at the Wellcome Sanger Institute.

## Authors contributions

LV and SP conceived the research. SP prepared the samples used for genome-wide analyses. PM performed bioinformatics analyses, analysed the data, design the website, and edited most of the figures. KJG performed differentiations in hPSCs using small molecule inhibitors and validations. RG, AO, DO and SB carried out co-immunoprecipitations and hPSC differentiation experiments and contributed to the project. PM and LV interpreted the data and wrote the manuscript. L.V. supervised, and supported the study, wrote and provided final approval of the manuscript.

## Data availability

All processed genome-wide datasets are publicly accessible in a genome browser at http://ngs.sanger.ac.uk/production/endoderm. Raw ChIP-seq data collected at 0h, 24h, 48h, and 72h is available at GEO DataSets (PRJNA593217). Raw ChIP-seq for 12h and 36h, ATAC-seq and RNA-seq data will be available in ArrayExpress upon publication under accessions E-MTAB-9276, E-MTAB-9124 and E-MTAB-9194, respectively.

## Code availability

Code used to analyse sequencing datasets is available at: https://github.com/pmb59/endoderm/

## Supplementary Information

### Cell culture of hESCs

hESCs (H9 from WiCell) were grown in defined culture conditions as described previously (Brons et al., 2007). H9 cells were passaged weekly using collagenase IV and maintained in chemically defined medium (CDM) supplemented with Activin A (10 ng/ml) and FGF2 (12 ng/ml). Pluripotent cells were maintained in Chemically Defined Media with BSA (CDM-BSA) supplemented with 10ng/ml recombinant human Activin A and 12ng/ml recombinant human FGF2 (both from Dr. Marko Hyvonen, Dept. of Biochemistry, University of Cambridge). Cells were passaged every 4-6 days with collagenase IV as clumps of 50-100 cells and dispensed at a density of 100-150 clumps/cm^2^. The culture media was replaced 48 hours after the split and then every 24 hours. Alternative culture conditions were used to maintain hPSCs used to study the role of ERK5-MAPK, p38-MAPK and JNK-MAPK. In sum, H9 and hIPSC lines FSPS13B and cA1ATD were routinely maintained on Vitronectin (StemCell Technologies)-coated plates in Essential 8 (E8) medium (Life technologies). Cells were passaged every 5-7 days using 0.5 uM EDTA and plated onto fresh vitronectin-coated plates in E8 medium. Medium was refreshed every day. This change corresponds to modification of protocols in our lab and has no influence on experimental outcomes.

### FUCCI-hESCs lines

The generation of FUCCI-hESC lines has been described in (Pauklin and Vallier 2013) and are based on the FUCCI system described in (Sakaue-Sawano et al., 2009).

### In vitro differentiation of hESCs

FUCCI-hESCs were differentiated into endoderm described previously (Vallier et al, 2009). Differentiation into endoderm was performed for up to 72 hours with a combination of cytokines as described in (Pauklin and Vallier, 2013; Pauklin et al. 2016). For cells sorted by FACS, the cells were collected and immediately placed into the endoderm differentiation media. Endoderm specification was performed in CDM with polyvinyl Alcohol (CDM-PVA) prepared without insulin and supplemented with 50ng/ml FGF2, 1µM Ly-294002 (Promega), 100ng/ml Activin A, and 10ng/ml BMP4 (R&D) for 3 days. Alternatively, and for cells grown in E8 medium, H9, FSPS13B or cA1ATD cells were plated as single cells onto gelatin/MEF-coated plates in E8 medium supplemented with 10uM Y-27632. The medium was refreshed the next day. Chemically Defined Media with Polyvinyl Alcohol (CDM-PVA) containing 100 ng/ml recombinant Activin A (CSCR, University of Cambridge), 80 ng/ml FGF2 (R&D Systems), 10 ng/ml BMP4 (CSCR, University of Cambridge), 10 μM LY29004 (Promega), and 3 μM CHIR99021 (Selleck Chemicals) was applied to the cells for 24 hours. The media was then replaced with fresh CDM-PVA supplemented with 100 ng/ml recombinant Activin A (CSCR, University of Cambridge), 80 ng/ml FGF2 (R&D Systems), 10 ng/ml BMP4 (CSCR, University of Cambridge) and 10 μM LY29004 (Promega). The next day, the media was removed and RPMI media supplemented with 1X B27 (Lifetech), 100 ng/ml Activin A, 80 ng/ml FGF2 and 1X non-essential amino acids (Lifetech) was added to the cells. To investigate the roles of ERK5-MAPK, p38-MAPK and JNK-MAPK, the endoderm differentiation media was supplemented with 10 µM U0126-EtOH, 10 µM SB203580 or 1 vM JNK-IN-8 respectively.

### Cell sorting by FACS

FACS on FUCCI-hESCs was performed as described before (Pauklin and Vallier, 2013; Sakaue-Sawano et al., 2009). In sum, hESCs were washed with PBS and detached from the plate by incubating them for 10 min at 37 °C in Cell Dissociation Buffer (Gibco). Cells were then washed with cold filter sterilised 1% BSA in PBS, before incubating cells in PBS 1% BSA with Tra-1-60 primary antibody (1:100) and Alexa Fluor 647 donkey α-mouse secondary antibody (1:1000) on ice for 20 min in the dark with occasional gentle mixing. The cells were then washed once with at least 50x pellet volume PBS 1% BSA, resuspended gently in 3ml sterile hESC maintenance media, and subjected to cell sorting by gating Tra-1-60+ cells according to the mAG/mKO2 FUCCI signals. The cell sorting was performed with a BeckmanCoulter MoFlo MLS high-speed cell sorter (Addenbrookes Hospital Flow Cytometry Core Facility) by using parameters described previously (Pauklin and Vallier, 2013), and the cells were sorted directly into collection tubes with 2 ml hESC maintenance media. After sorting, the cells were pelleted and placed in endoderm differentiation media. The media was changed every 24 hours and samples were collected at different timepoints for subsequent processing for RNA-Sequencing, histone ChIP-Sequencing and ATAC-Sequencing.

### Immunostaining

The immunostaining method has been described previously (Pauklin et al., 2013; Pauklin et al., 2016; Bertero et al., 2015). Cells were fixed for 20 minutes at 4°C in PBS 4% PFA (electron microscopy grade), rinsed three times with PBS, and blocked and permeabilized at the same time for 30 minutes at room temperature using PBS with 10% Donkey Serum (Biorad) and 0.1% Triton X-100 (Sigma). Incubation with primary antibodies (**Table S2**) diluted in PBS 1% Donkey Serum 0.1% Triton X-100 was performed overnight at 4°C. Samples were washed three times with PBS, and then incubated with AlexaFluor secondary antibodies (**Table S2**) for 1 hour at room temperature protected from light. Cells were finally washed three times with PBS, and Hoechst (Sigma) was added to the first wash to stain nuclei. Images were acquired using a LSM 700 confocal microscope (Leica).

### Flow-cytometry

Single cell suspensions were prepared by incubation in Cell Dissociation Buffer (Gibco) for 10 minutes at 37° followed by gentle pipetting. Cells were fixed in 4% PFA for 20 min at 4°C. This was followed by permeabilization and blocking with 10% serum + 0.1% Triton X-100 in PBS for 30 min at RT and incubation with primary antibody in 1% serum + 0.1% Triton X-100 for 2h at 4°C. After washing the samples three times with PBS, they were incubated with a secondary antibody for 2h at 4°C, washed three times with PBS and analysed by flow cytometry. Flow-cytometry was performed using a Cyan ADP flow-cytometer and at least 20,000 events were recorded. Data was analysed by FloJo software. Cell cycle distribution was analysed by Click-It EdU incorporation Kit (Invitrogen) according to manufacturer’s guidelines.

### RNA-seq experiments

Samples for RNA-sequencing were collected at different time points (0, 12, 24, 36, 48, 60, 72h) from FUCCI-hESCs differentiated to endoderm. The libraries for RNA-sequencing were generated by the Wellcome Sanger Institute Illumina Bespoke Sequencing Facility and sequencing was performed onsite. The libraries were generated at a library fragment size between 100 bp to 1 kb with Stranded RNAseq Standard with Oligo dT pulldown. The samples were multiplexed and analysed by Illumina Hiseq V4 with a paired end read length PE75. All samples were amplified with a standard 10 PCR cycle before sequencing. Samples were distributed equally across sequencing lanes and a total of 2,037,382,345 mapped reads with MAPQ ≥ 10 were obtained after sequencing using Illumina HiSeq 2000 (97,018,207 reads/sample on average). Processed RNA-seq data is freely available at http://ngs.sanger.ac.uk/production/endoderm.

### ChIP-seq experiments

ChIP-seq was performed using FUCCI-Human Embryonic Stem Cells (FUCCI-hESCs, H9 from WiCell) in a modified endoderm differentiation protocol (see details below). Cells were grown in defined culture conditions as described previously (Brons et al 2017). Pluripotent cells were maintained in Chemically Defined Media with BSA (CDM-BSA) supplemented with 10 ng/mL recombinant Activin A and 12 ng/mL recombinant FGF2 (both from Dr. Marko Hyvonen, Dept. of Biochemistry, University of Cambridge) on 0.1% Gelatin and MEF media coated plates. Cells were passaged every 4–6 days with collagenase IV as clumps of 50–100 cells. The culture media was replaced 48 h after the split and then every 24 h. The generation of FUCCI-hESC lines is based on the FUCCI system (Pauklin, S. & Vallier, 2014; Sakaue-Sawano, A. et al. 2008). hESCs were differentiated into endoderm as previously described (Vallier et al, 2009). Following FACS sorting, Early G1 (EG1) cells were collected and immediately placed into the endoderm differentiation media and time-points were collected at 0h, 12h, 24h, 36h, 48h and 72 h. Endoderm specification was performed in CDM with Polyvynilic acid (CDM-PVA) supplemented with 20 ng/mL FGF2, 10 µM Ly-294002 (Promega), 100 ng/mL Activin A, and 10 ng/mL BMP4 (R&D). We performed ChIP-sequencing for various histone marks (H3K4me3, H3K27me3, H3K4me1, H3K27ac, H3K36me3) (see **Supplementary Table 1** for antibodies), on two biological replicates per condition (Pauklin et al, 2016), except at 36h time-point, where only one replicate was obtained, and we could not generate H3K27me3 (H3K27me3 for 36h sample failed, but we do expect little changes based in comparison 24-48h) At the end of the ChIP protocol, fragments between 100 bp and 400 bp were used to prepare barcoded sequencing libraries. 10 ng of input material for each condition were also used for library preparation and later used as a control during peak calls. The libraries were generated by performing 8 PCR cycles for all samples. Equimolar amounts of each library were pooled, and this multiplexed library was diluted to 8pM before sequencing using an Illumina HiSeq 2000 with 75 bp paired-end reads. Processed ChIP-Seq data is freely available at http://ngs.sanger.ac.uk/production/endoderm.

### ATAC-Seq experiments

hESCs were sorted to early G1 phase and plated at 200,000 cells per well in 12-well plates with 0.5ml endoderm differentiation media. After differentiating the cells for a range of time points, the cells were washed once with PBS, collected in Cell Dissociation Buffer (Gibco 13150-016) and centrifuged at 300g for 3 min. The cell pellets were then resuspended in 2 ml of 4°C PBS and counted by haemocytometer for using 100,000 cells in the subsequent step. Cells were centrifuged at 300g for 3 min, the supernatant aspirated, the cell pellet resuspended in 150 ul of Isotonic Lysis Buffer (10 mM Tris-HCl pH 7.5, 3 mM CaCl, 2 mM MgCl2, 0.32 M Sucrose and Protease Inhibitors, Roche), and incubated for 12 min on ice. Triton X-100 from a 10% stock was then added at a final concentration of 0.5%, the samples were vortexed briefly and incubated on ice for 6 min. The samples were centrifuged for 5 min at 400g at 4°C, and the cytoplasmic fraction removed from the nuclear pellet. The samples were resuspended gently in 1 ml of Isotonic Lysis Buffer and transferred to a fresh 1.5 ml eppendorf tube. The nuclei were centrifuged at 1500g for 3 min at 4°C and the supernatant aspirated thoroughly from the nuclear pellet. This step was immediately followed by tagmentation (Nextera DNA Sample Preparation Kit for 24 Samples, FC-121-1030) by resuspending each sample in 50 µl Nextera mastermix (25 µl TD buffer, 20 µl of water and 5 µl of TDE1 per reaction). The nuclear pellet was resuspended thoroughly by pipetting and incubated at 37 °C for 30 mins. The reaction was stopped with 250 µL of buffer PB from the Qiagen PCR purification kit, followed by Qiagen PCR clean up protocol using MinElute columns and eluting each sample in 11.5 µl buffer EB. For the control sample, the nuclear pellet was subjected to genomic DNA isolation with GenElute Mammalian Genomic DNA Miniprep Kit (Sigma, G1N70) according to manufacturer’s protocol, and the purified genomic DNA was thereafter immediately used for tagmentation as for other ATAC-seq samples. Next a PCR reaction (for all samples including control sample) was performed with the following constituents: 10 µl template from tagmentation, 2.5 µl I7 primer (Nextera® Index Kit with 24 Indices for 96 Samples, FC-121-1011), 2.5 µl I5 primer, 2.5 ul Nextera cocktail and 7.5µl Nextera PCR mastermix. The PCR settings were as follows: initial denaturation at 98 °C for 30 seconds, then 12 cycles of 98 °C for 10 seconds, primer annealing at 63 °C for 30 seconds and elongation at 72 °C for 3 minutes, which was followed by a final elongation at 72 °C for 3 minutes and holding at 10 °C. After completing the PCR, the sample volumes were increased to 50 µl by adding Qiagen EB buffer from the PCR purification kit. The PCR primers were removed with 1 x 0.9:1 SPRI beads (Beckman Coulter, Cat no. A63880) according to manufacturer’s protocol and samples eluted in 20 µl. DNA size-selection was performed as follows: the samples were run on a 1 % agarose gel in TAE Buffer at 90 V for 25 minutes. The DNA was cut within the range of 150 bp to 1 kb and purified by Qiagen Gel Extraction Kit with MinElute columns, by eluting in 20 µL Qiagen buffer EB. 1 µl of the samples were run on Agilent HS Bioanalyzer HS for confirming the size selection of the ATAC libraries. ATAC-sequencing was performed by Illumina HiSeq 2000 sequencing with 75 bp PE for obtaining 460,130,600 million reads per library on average (5,981,697,796 in total for 0, 12, 24, 36, 48, 72h samples in duplicates plus a control sample). Processed ATAC-seq data is freely available at http://ngs.sanger.ac.uk/production/endoderm.

### Quantitative real-time PCR

Media was removed and cells were washed once with DPBS (Life technologies) before 300 ul of RNA lysis buffer was added. RNA was extracted using the GenElute Mammalian Total RNA Miniprep Kit (Sigma-Aldrich) per manufacturer’s instructions. 500 ng RNA was reverse-transcrived using random primers (Promega), dNTPs (Promega), RNAseOUT (Invitrogen), DTT (Invitrogen) and SuperScript II (Invitrogen). QPCR reactions were made up using 2x KAPA SYBR Fast qPCR Master Mix kit (Kapa Biosystens), 4.2 ul of 30x diluted cDNA, and 200 nM of forward and reverse primers. Samples were run in the QuantStudio 12K Flex real-time PCR system machine and a 384-well plate and analysed using the delta-delta cycle threshold (Ct) method normalized to housekeeping gene, ACTB. Primer sequences can be found in Supplementary Table S3.

### RNA-Seq data analysis

Reads were mapped to the human genome (GRCh38.15) using TopHat v2.0.13 (Kim et al., 2013) with the following options: “--library-type fr-firststrand”, “--mate-inner-dist 100 --no-coverage-search --microexon-search” and “—transcriptome-index” with a TopHat transcript index built from ensembl_76_transcriptome-GRCh38_15.gtf. Reads with Mapping Quality Values <10 were filtered out with samtools. featureCounts was used on paired-end reads to count fragments in annotated gene features, with parameters ‘-p -C -T 8 -t exon -g gene_id’ (Liao et al., 2014). DESeq2 R/Bioconductor package was used in differential gene expression analysis between samples, requiring at least a 1.5-fold expression change and a Benjamini-Hochberg adjusted *P*-value smaller than 0.01 (Love et al., 2014) for a gene to be deemed as differentially expressed. The function ‘rpkm’ in the R/Bioconductor package edgeR (Robinson et al., 2010) was used with default parameters to normalize count gene expression. Raw bedGraphs were normalized per million mapped reads in the library per library size in all samples (Conesa et al., 2016). Spearman’s correlation *ρ* values were calculated in R for FPKM expression values of genes expressed at more than 5 FPKM in at least one of the samples under comparison. Hierarchical clustering of *ρ* values clustered all the triplicates in each condition together. (Extended Data Fig. 2b). PCA implemented in DESeq2 was performed (Extended Data Fig. 2c). Other bioinformatics analyses were carried out following standard procedures (Conesa et al 2016).

### Heatmap of gene expression in Fig.1

Average of FPKM values was computed for all the replicates of each condition. The function ‘Heatmap’ in the package ‘ComplexHeatmap’ was used to perform row clustering in selected group of genes using Euclidean distance and ‘complete’ clustering method. For the heatmap in Supp. Figure 1d scaled log2(FPKM+1) was used for all differentially expressed, for ‘protein_coding’ biotypes.

### K-means clustering

Model-based optimal number of clusters *K* that minimized Bayesian Information Criterion (BIC) was considered for 6,317 differentially expressed protein-coding genes. The number of clusters *K* = 13 was selected as the one that minimized the BIC using the function ‘mclust’ in the R package ‘Mclust’. To smooth the data for representing the curves in Supp. Fig. 1e, we used functional data analysis R package “fda” v2.4.4. First, we represented data values using 5 B-spline basis functions located a 0, 12, 24, 36, 48, 60, 72h without roughness penalties in the second derivative (*λ* = 0). We used the functions create.bspline.basis() and smooth.fd() over the interval 0–72 h. Then we evaluated the mean and the s.d. of the functional data objects using the R functions ‘mean.fd’ and ‘sd.fd’.

### ChIP-seq data analysis

#### Preprocessing and peak calling

Reads were mapped to GRCh38 reference assembly using BWA (Li and Durbin, 2009). Only reads with mapping quality score ≥10 and aligned to autosomal and sex chromosomes were kept for further processing. Peak calling analysis (Bailey et al, 2013) was performed using PeakRanger (Feng et al, 2011), and only the peaks that were reproducible at an *FDR* of ≤0.05 in two biological replicates were selected for further processing. Peak calling was done using appropriate controls with the tool peakranger 1.18 in modes *ranger* (H3K4me3, H3K27ac; ‘-l 316 -b 200 -q 0.05’), *ccat* (H3K27me3; ‘-l 316 --win_size 1000 --win_step 100 --min_count 70 --min_score 7 -q 0.05’) and *bcp* (H3K4me1, H3K36me3; ‘-l 316’). Adjacent peak regions closer than 40 bp were merged using the BEDTools suite (Quinlan and Hall, 2010), and those overlapping ENCODE blacklisted regions were filtered out (ENCODE Excludable Mappability Regions; ENCODE Project Consortium et al, 2012). bedGraph format files were produced for each sample using BEDTools 2.17.0 (Quinlan and Hall 2010). The reads mapped at both DNA strands from 5’ to 3’ direction were extended to a length of 316 bp, and the readcount at each genomic position was normalized to the library size and per million reads (multiplying every value by ‘1,000,000 / number_of_mapped_reads’). bedGraph files were converted to bigWig using UCSC tool bedGraphToBigWig, and are available for visualisation on the Biodalliance genome viewer (Down et al. 2011) at: http://ngs.sanger.ac.uk/production/endoderm.

#### Differential peak calling in histone modification ChIP-seq

G-tests implemented in diffReps (Shen et al. 2013) were used to detect differential histone modification regions. hg38 as reference genome, and an average fragment size as calculated previously (rest of parameters default). Because we have only 1 replicate at 36h, we decided to use G-tests instead of Negative Binomial tests for all the comparisons, as recommended in Shen et al. 2013. All input samples were merged and used as background control. Differential histone modifications regions not overlapping (at least 1bp) significant chromatin marks previously detected during peak calling at least in one of conditions under comparison were removed. Regions were ranked by their adjusted *P*-value and reported as differentially enriched only if the absolute *FC*≥2, and Benjamini-Hochberg corrected *P-*value ≤ 0.01. Genes in a 10 kb window of the regions were reported.

| Dataset | --mode | --window | --step | --nsd | --frag | --gap | --pval |
| --- | --- | --- | --- | --- | --- | --- | --- |
| H3K4me3 | peak | 600 | 60 | 20 | 316 | 30 | 0.000001 |
| H3K27ac | peak | 600 | 60 | 20 | 316 | 100 | 0.000001 |
| H3K27me3 | block | 2000 | 200 | 10 | 316 | 50 | 0.000001 |
| H3K4me1 | block | 6000 | 300 | 5 | 316 | 200 | 0.000001 |
| H3K36me3 | block | 10000 | 200 | 2 | 316 | 200 | 0.000001 |

### ATAC-seq data analysis

#### Preprocessing and peak calling

A total of 5,981,697,796 PE reads (75 bp) were sequenced using Illumina HiSeq 2000, which includes one deeply sequenced control sample of 369,590,751 reads. BWA v0.7.12 (Li and Durbin, 2010) with parameters ‘mem -t 16 -p -T 0’ was used for read alignment against hg38 (GRCh38.15) reference assembly of the human genome. Aligned reads were retained if MAPQ ≥ 5. Only autosomal and sex chromosomes were retained. Mitochondrial contamination (% reads) was found proportional to the differentiation stage (high (∼80%) in hESCs and low in Definitive Endoderm (∼25%)). Peak calling against the control sample was performed using JAMM 1.0.7.1 (Joint Analysis of NGS replicates via Mixture Model clustering), which proved to improve accurate determination of peak boundaries (Ibrahim et al., 2014), with parameters ‘-m normal -r region -f 1,1,1 –b 140’. Duplicates were removed at this stage as in Buenrostro et al. (2013) to improve peak calling, same as recommended in ChIP-seq data analysis (Bailey et al., 2013). To select the number *n* of reproducible peaks at IDR ≤ 0.05, Irreproducible Discovery Rate (IDR) analysis (Li et al., 2011) was performed on JAMM’s ‘filtered’ peaks of individual replicates using JAMM’s peak scores *S*_*p*_, which are defined as:

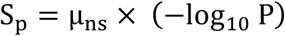

where *μ*_*ns*_ is the mean peak background normalized signal, and *P* is the Benjamini-Hochberg corrected *P*-value of Mann-Whitney-U non-parametric tests. A number of peaks equal to the minimum number of peaks in one of the replicates were submitted for IDR analysis. Top *n* peaks in JAMM’s replicate integration (pooled replicates) method were then selected as the highly confident set. Normalised signal tracks were built after extending each read to estimated average fragment size of 140bp. We generated a consensus-merged list of 253,618 peak regions, filtered out those overlapping the human ENCODE blacklisted genomic regions (https://sites.google.com/site/anshulkundaje/projects/blacklists), and considered a final set of 253,349 open chromatin regions for further analyses.

#### Differential open chromatin region analysis

To statistically identify regions of differential open chromatin in ATAC assays, a modified version of function narrowpeaksDiff.R in the Bioconductor package NarrowPeaks (Mateos et al., 2015) was used. We performed Hotelling’s *T*^2^ tests on the functional principal component scores to identify significant differences across conditions. Genomic regions were declared significant if Bonferroni adjusted *P*-value ≤ 10^−4^ and absolute Fold-Change (FC) ≥ 2.0 in a set of aggregated regions of those under comparison. 31 bins were used for signal extraction in the R package ‘genomation’ and 10 equidistant B-splines bases were used for functional principal component analysis (FPCA, first FPC considered). Regions were then ranked then by the scores:

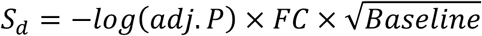

(**data available here: http://ngs.sanger.ac.uk/production/endoderm/**), where baseline is defined as ½*(Avg. normalized signal in time-point 1+Avg. normalized signals in time-point 2). diffNGS source code is available at http://github.com/pmb59/diffNGS. Genes were associated to peaks if the former were located in a region 10kb upstream or downstream from differential open chromatin regions. Open chromatin regions with no change at any stage were classified as ‘invariant’.

#### Motif analysis of DNA sequences in differential ATAC-seq regions

Motif enrichment analysis was performed with HOMER2 (v4.8.3), using the function ‘scrambleFasta.pl’ to create a set of background frequencies (with the same number of sequences, as recommended). We scored known motifs for enrichment in the FASTA files using the function ‘known’, and motifs from the CIS-BP database (3,059 PWMs; http://cisbp.ccbr.utoronto.ca; Weirauch et al., 2014) - those were used also for footprinting analysis (see below). Occurrences were ranked and a scatter plot was generated using their Log(*P*-value) and Log2(*Enrichment Ratio*) for TF motifs associated to genes expressed at least 1.0 FPKM (mean across triplicates) in the earliest time point of the 2 consecutives being analysed. TF_Name was associated to PWMs using the CISBP information file TF_Information.txt (of directly determined motifs and best inferred motifs).

#### Pearson correlation between RNA-seq and ATAC-seq

Pearson’s product-moment correlation was calculated in R using the function cor.test (alternative hypothesis: true correlation is not equal to 0). Log_2_ Fold-changes obtained in DESeq2 for protein-coding genes were used. To obtain the Fold change in ATAC-seq, maxima of normalized ATAC-seq signal in a region 10kb upstream the promoter of protein-coding genes was obtained with ScoreMatrixBin (genomation package) in each ATAC-seq replicate of consecutive time points, and log2 of the fold change was computed.

#### GO enrichment analysis in variant open chromatin regions

Ensembl gene IDs of protein_coding genes were converted to Entrez gene IDs using biomaRt (ensemble.76). Gene Ontology (GO) enrichment analysis was performed using GOstats (Falcon and Gentleman, 2007), Hypergeometric Tests for GO term association (function ‘hyperGTest’) were run with BP ontology and *P*-value cutoff = 0.001. This function computes Hypergeometric *P*-values for over- (or under-) representation of each GO term in the specified ontology among the GO annotations for the genes of interest.

#### Identification of footprints of DNA-binding transcription factors

We used FootPrintMixture (Yardicimci et al., 2014) to detect footprints of TFs in ATAC-seq, adapting this tool to explicitly control for assay-specific sequence bias, which is fundamental for DNase-seq and ATAC-seq data analysis (Sung et al., 2014; Madrigal, 2015). PWMs of “Directly determined or best inferred motif” for Homo sapiens were downloaded from CIS-BP database (Weirauch et al., 2014).

#### Controlling the sequence-bias with a deproteinized ATAC control

We retrieved Tn5 transposition sites from the deproteinized ATAC control sample (328,759,008 mapped reads) after translating 5’-ends of the reads +4 bp for insertion on forward strand, and -5 bp in the reverse strand (Buenrostro et al. 2013). Then we obtained 6-mers frequencies ±3 bp around the transposition site for chr1 using Jellyfish (Marçais and Kingsford, 2011) with parameters “count -m 6 -s 4000M -t 4”. Background 6-mer counts in the human genome were estimated using the ‘fasta-get-markov’ program in the MEME Suite with a Markov Model of order *m* = 5 (we consider all genome to be mappable). Then, the values in SeqBias.txt in FootprintMixture would be *K*-mer frequencies in the deproteinized sample divided by background frequencies. Here deviations from 1.0 would indicate bias. It is likely that no substantial differences between Tn5 insertion preferences between purified genomic DNA and human chromatin, suggesting that the local insertion preference into chromatin is identical to that found in naked genomic DNA (Buenrostro et al. 2013; Madrigal 2015).

CisBP database of human 3,059 valid PWMs was used (Weirauch et al. 2014). Matches of the PWMs were obtained using FIMO (Grant et al, 2011) with a cut-off of *P*<10^−4^, then ranked, and motifs with at least 10k matches (up to a maximum to 500k matches) were submitted for 2-component mixture model in FootPrintMixture to infer TF binding ±25 bp around each motif. Extended motif occurrences that lie out of genomic regions in hg38 were not submitted to analysis. To get high quality reproducible footprints, motif matches with Footprint Likelihood Ratio (FLR) ≥10.0 in each biological replicate were classified as “Bound”. TF footprints that were present in both replicates, and with at least 50 FPs for the PWM, were considered as predicted TF BSs. FLR mean of the replicates was calculated. For this calculation, only unique genomic regions were taken into account (for those palindromic sequences with a footprint reported in each strand, we obtained the mean FLR). Normalized average of Tn5 insertion densities for ‘Bound’ sites were plotted ±100 bp around motif using R and NucleoATAC (Schep et al 2015) function ‘pyatac ins’ for whole genome, after normalizing for number of reads in the library. For visualization purposes reads in both replicates were merged.

Next, we computed the protection scores for footprint predictions of a PWM as proposed in Gusmao et al. 2016. This score is important because measures the quality for a given TF (PWM) and set of predictions (the higher, the most reliable the footprints are). Low protection scores will indicate TFs with short residency times in the DNA (like SOX2).

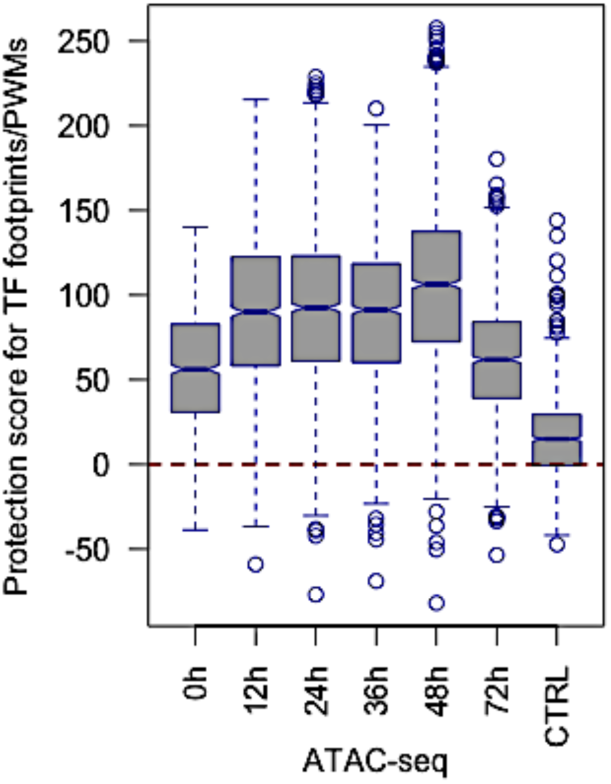

Flanking regions were considered ±25 bp upstream/downstream of the motif for this calculation [see red arrows in footprint plots in the Figures above].

Footprints with short DNA residency time are poorly detected and likely to be false positives. TFs with intermediate and long residency times have positive PS (Gusmao et al., 2016). Footprints with a negative protection score and associated to genes with ≤ 1.0 FPKM (no expression) were removed. PWMs associated to >1 TF were not considered.

We repeated the analyses in the control sample (considering the same sample as duplicate), without applying gene expression cut-off (0.0 FPKM). This demonstrates that ATAC-seq, as DNase-seq footprints, cleavage signatures are dictated by sequenced biases (Sung et al, 2014), and that filtering by Protection score (PS, red arrows in the figure below) is very recommended step in footprint determination.

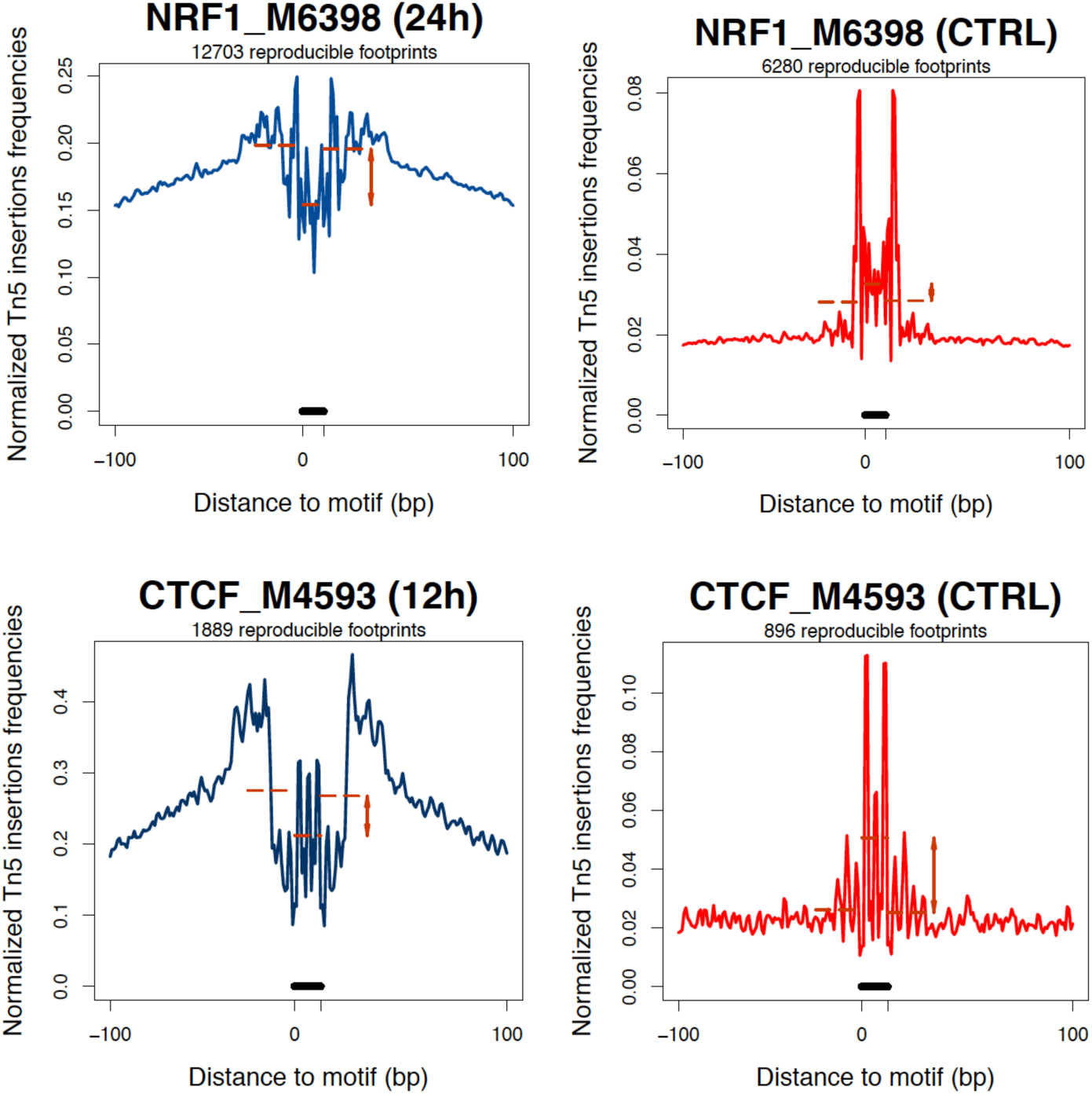

**Normalized Tn5 insertion frequencies for CiSBP PWMs M6398 (NRF1) and M4593 (CTCF) for ATAC-seq and control samples illustrate the importance of correcting Tn5 sequence bias**.

#### Motif-agnostic footprint detecting with Wellington

We run also Wellington/pyDNase 0.2.4 for digital genomic footprinting, a non-motif centric footprint algorithm detector (Piper et al. 2013) to search TF footprints between 4-30 bp (‘-A -fdr 0.1 --FDR_limit -4 --pv_cutoffs -4 -fp 4,30,1 -fdriter 500 -- one_dimension’). As search space we used the set of consensus open chromatin regions extended 50 bp upstream and downstream. BAM files of alignment for replicates were merged to achieve a better coverage. A relaxed set of footprints was obtained at each time point (*P* ≤ 1e-4).

#### Final set of TF footprints

Nonredundant footprints with some overlap with Wellington footprints were kept as the final set of curated footprints. In summary, the final set of curated nonredundant footprints were considered as as those:

- With FLR ≥10.0 (Footprint-Mixture), reproducible in both replicates.
- With Protection Score PS ≥ 0.
- With CiSBP PWM associated to only 1 TF.
- Expression of gene/TF ≥ 1.0 FPKM (from RNA-seq at the same stage).
- Only footprints with highest FLR were kept for Overlapping footprints (distinct PWMs) for the same TF (in case of equal FLR, take the 1^st^) to get nonredundant set.
- Finally, only nonredundant footprints detected also by Wellington with a relaxed cut-off were kept.

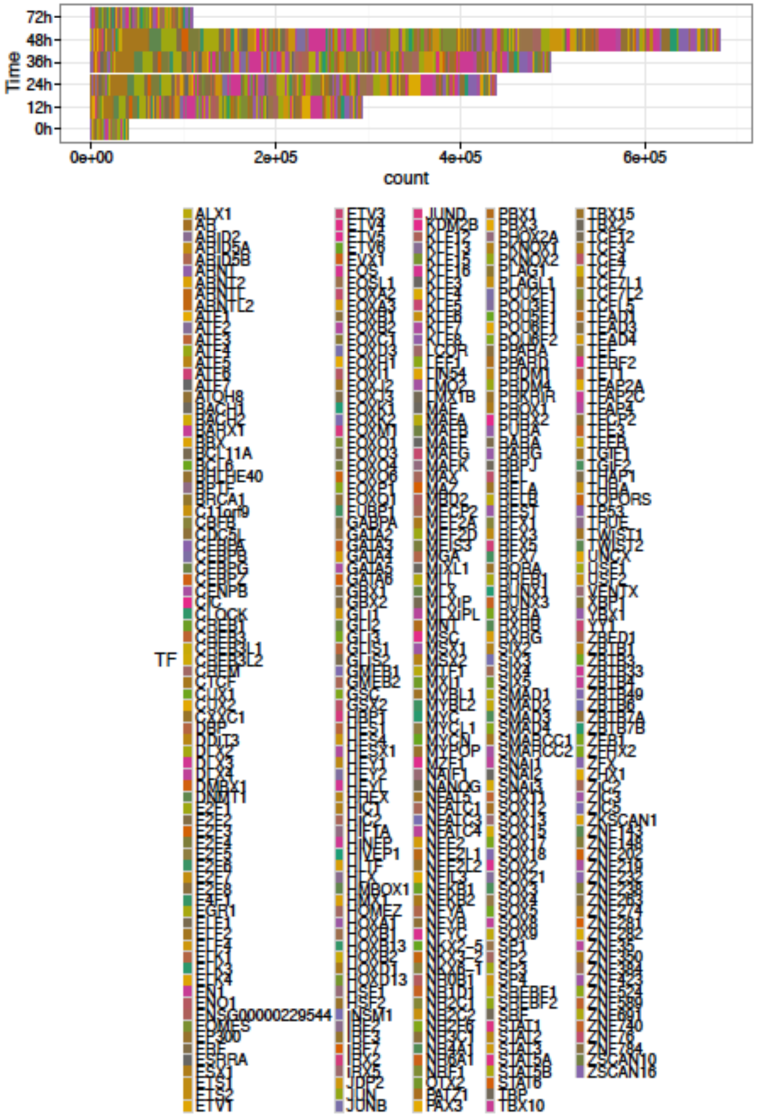

**Number of curated nonredundant final footprints detected during endoderm differentiation. Number of footprints is, partly, a consequence of sequencing depth**

### Overlap between H9-hESCs-EG1 and H7-hESCs ENCODE footprints

H7-hESCs ENCODE footprints (GRCh37/hg19) were downloaded from ftp://ftp.ebi.ac.uk/pub/databases/ensembl/encode/supplementary/integration_data_jan2011/byDataType/footprints/jan2011/. ENCODE footprints were converted to hg38 assembly using UCSC tool liftOver and the overlap was computed with BEDtools.

### Differential genomic footprinting with Wellington_bootstrap

pyDNase-0.2.4 was used to obtain differential footprints between consecutive time stages (http://pythonhosted.org/pyDNase). As a search space, a BED file with genomic locations was provided and consisted of the consensus open chromatin sites extended 50 bp each side. BAM files of alignment for replicates were merged and evaluated with Wellington_bootstrap with parameters “-A -fp 4,30,1 -fdr 0.05 -fdrlimit -10 -fdriter 100”. Importantly, Wellington_bootstrap allows to control for different sequencing depth in the two samples (Piper et al., 2015). A score *S*≥5 was used for differential footprints to be considered as such. The overenrichment of the number of predictions for a TF was statistically assessed considering the proportion over the total footprints against its relative proportion in the differential list by applying Chi-squared tests. Benjamini–Hochberg (BH) multiple hypothesis testing correction was applied.

### Global TF footprint pattern change between conditions

Bivariate genomic footprinting was performed with BaGFoot v.0.9.7 (Baek et al, 2017) over motifs of human transcription factors from the CiSBP database (http://cisbp.ccbr.utoronto.ca).

### Peak annotation in ATAC-seq peaks and TF footprints

NIH PAVIS v.04-08-2016 was used for peak-to-gene annotation (GRCh38r76/hg38) of differential ATAC-seq peaks (https://manticore.niehs.nih.gov/pavis2/). Upstream Length: 10kb, Downstream Length: 5kb (rest of parameters under default configuration). TF footprints unique regions were obtained using the function ‘merge’ in BEDTools before submitting to PAVIS.

### Functional linear regression with a scalar response

Functional Data Analysis (FDA) has been applied in genomics to allow data analysis on continuous data (Cremona et al., 2019). Using a functional linear model (Ramsay and Silverman, 2005) instead of multivariate linear model allows including in the model the spatial dependency between TF binding and chromatin change. The model used to predict chromatin accessibility change exerted by footprints of a transcription factor was:

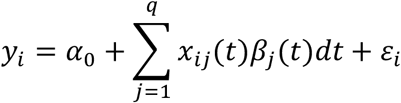

where *y*_*i*_ is the chromatin change of region *i* between time-point *t* and time-point *t*+1, *x*_*i*_*(t)* is a functional covariate (q=1) defined by the fooptrinting signal, *α*_0_ is the intercept term, *β*_*j*_ are the regression coefficient functions, and *ε*_*i*_ is the independent and identically distributed (i.i.d.) error term. Only differential accessibility regions with a Fold-change *FC*> 1.5 were considered. *FC* ≥ |10| were removed and not considered to avoid outliers and 1000 bp were used around the centre of each ATAC-seq peak. Log_2_ (abs(*FC*)) was considered as the scalar variable *y*_*i*_ to be regressed. A maximum of 50,000 genomic regions, ranked by their abs FC, were considered in the model. Putative TF binding sites were represented in the functional covariate as 1.0 (absence as 0.0) using five order-4 B-splines. The linear model was only considered when at least 15 footprints overlapped regions of interest, and only TF with footprints in >50 of the selected regions were implemented in the model. Squared multiple correlation (*R*^*2*^) and *F*-ratio were used to assess the improvement of fit. *P*-values were computed for the *F*-statistic using the ‘pf’ function in R. The R package ‘fda’ was used in these analyses.

### Jaccard Index overlap analysis and clustering

We used the R package GenometriCorr 1.1.17 (http://genometricorr.sourceforge.net/) (Favorov et al., 2012) to calculate the Jaccard Index (*JI*), a measure of correlation between two genomic intervals, for CHIP-seq and ATAC-seq peaks and differential sites (up and down, only those >100 in number). Clustering was performed with ‘Heatmap’ function of the Bioconductor package ComplexHeatmap.

**Supplemental Table S2.**
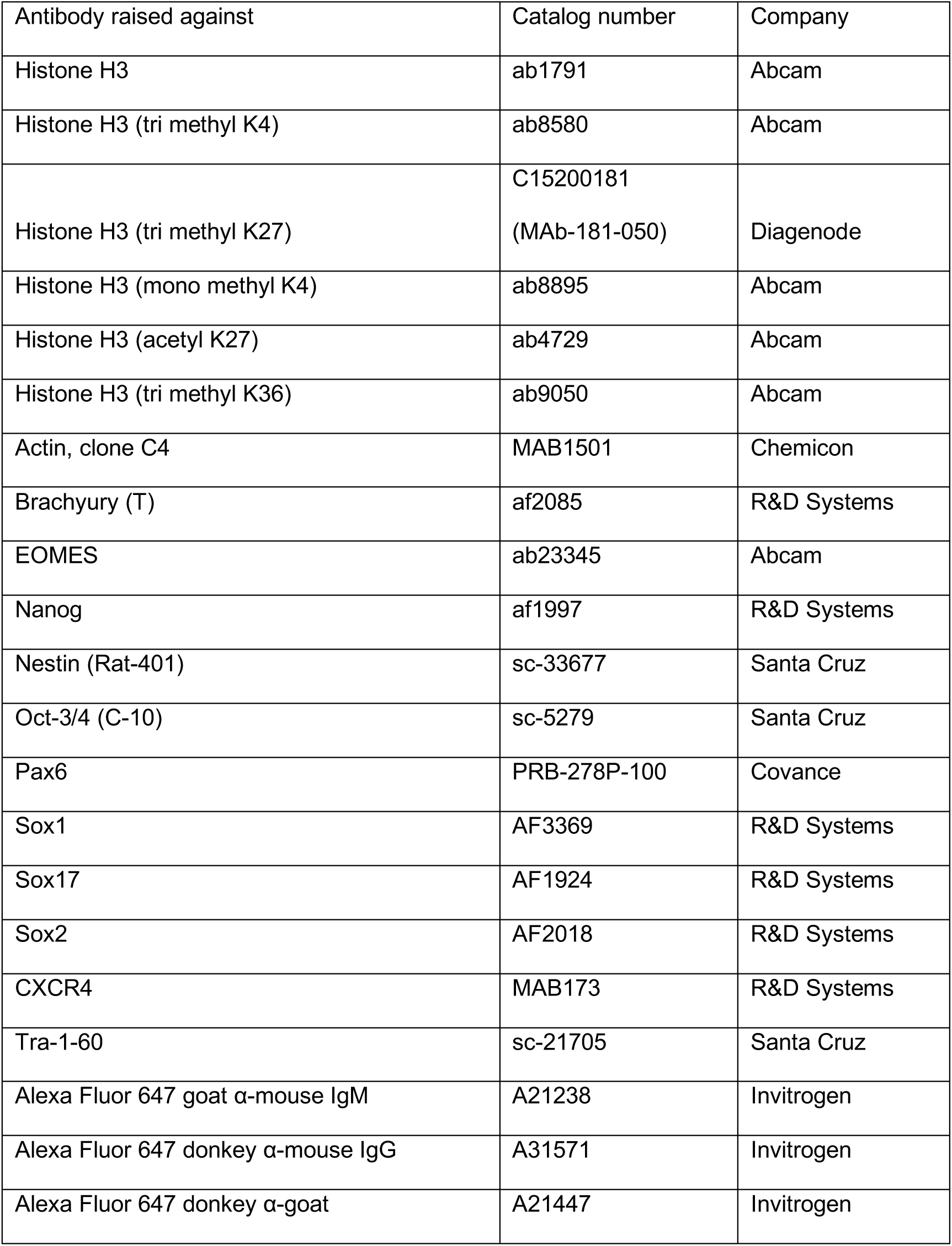
Antibodies.

**Supplemental Table S3.**
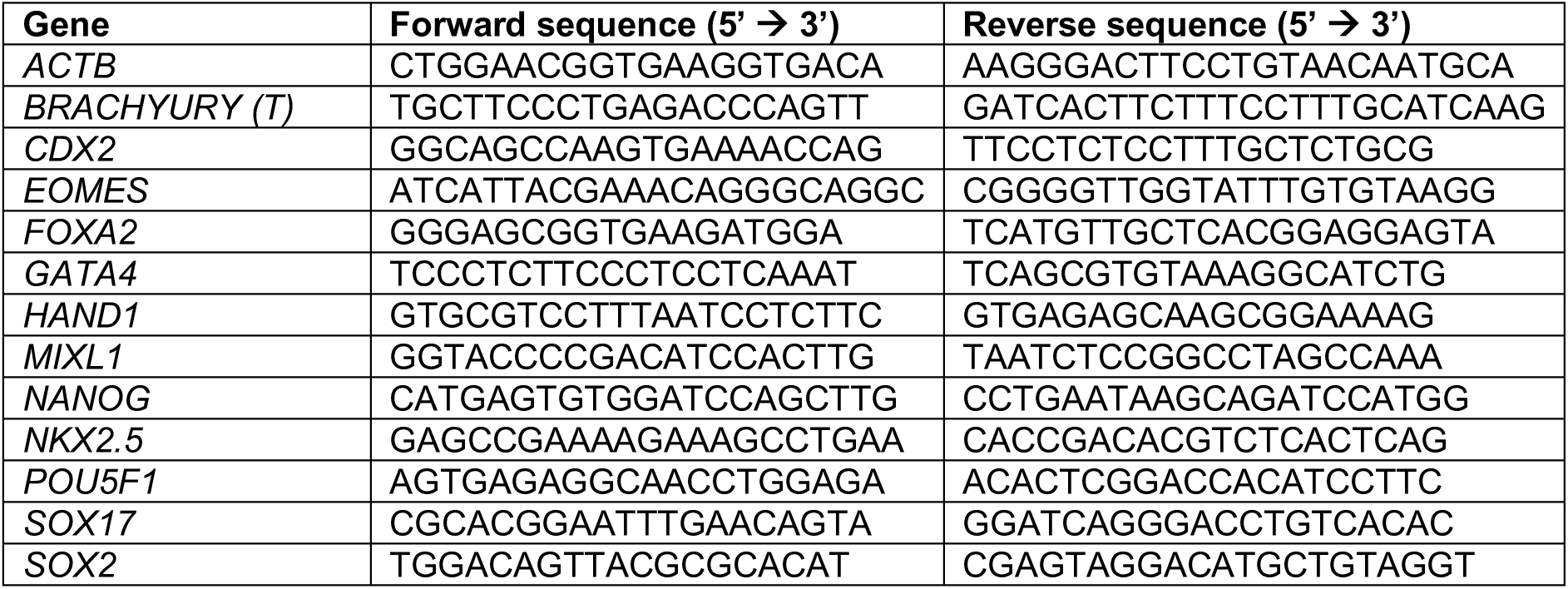
QPCR Primers.

## Supplementary Figure Legends

**Supplementary Table 1:**
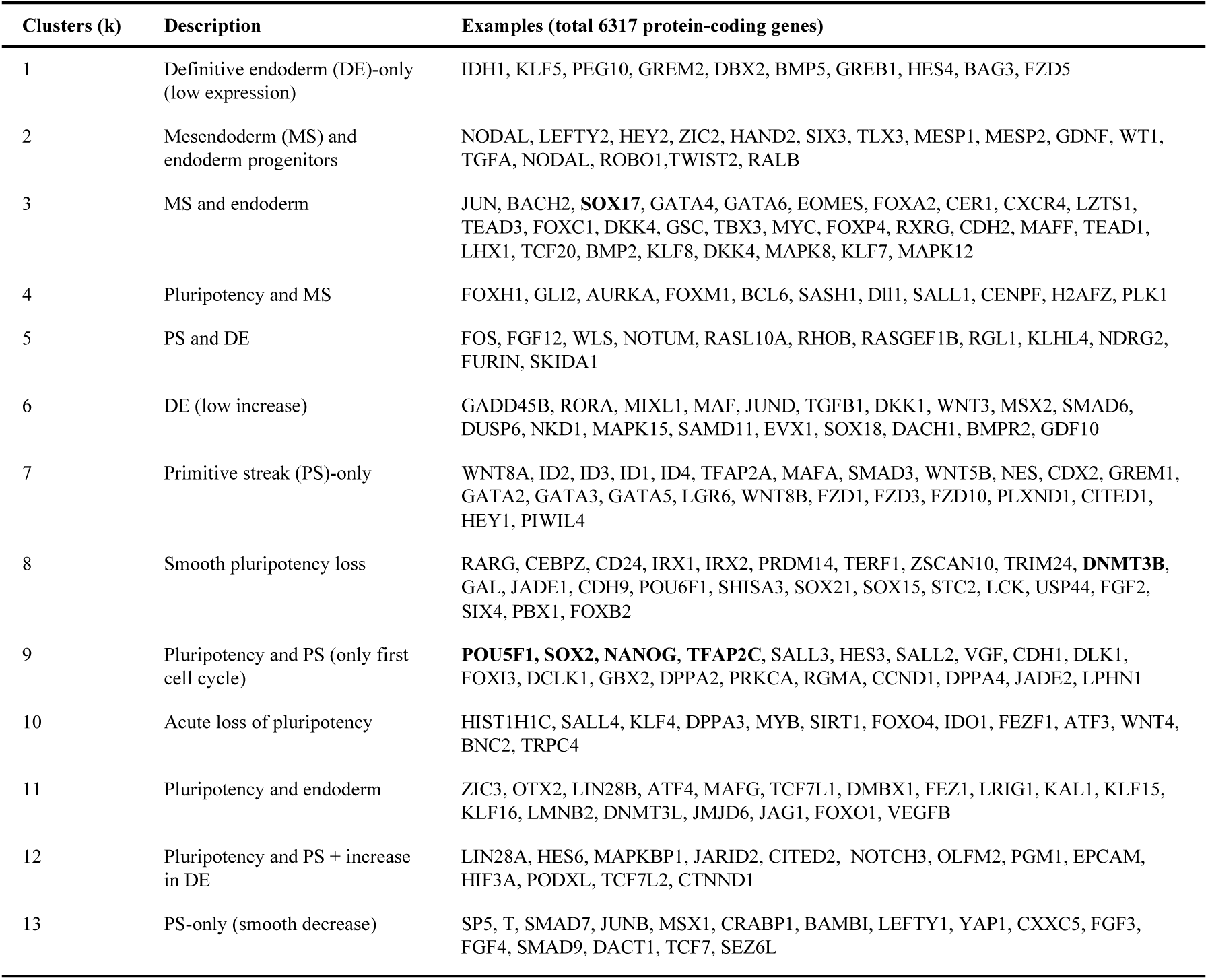
Genes expressed in the different clusters identified by k-means clustering of differentially expressed genes during differentiation of EG1 FUCCI-hPSCs (0h, 12h, 24h, 36h, 48h, 60h and 72h) as denoted in Fig. S1d and S1e.

**Supplementary Figure 1:**
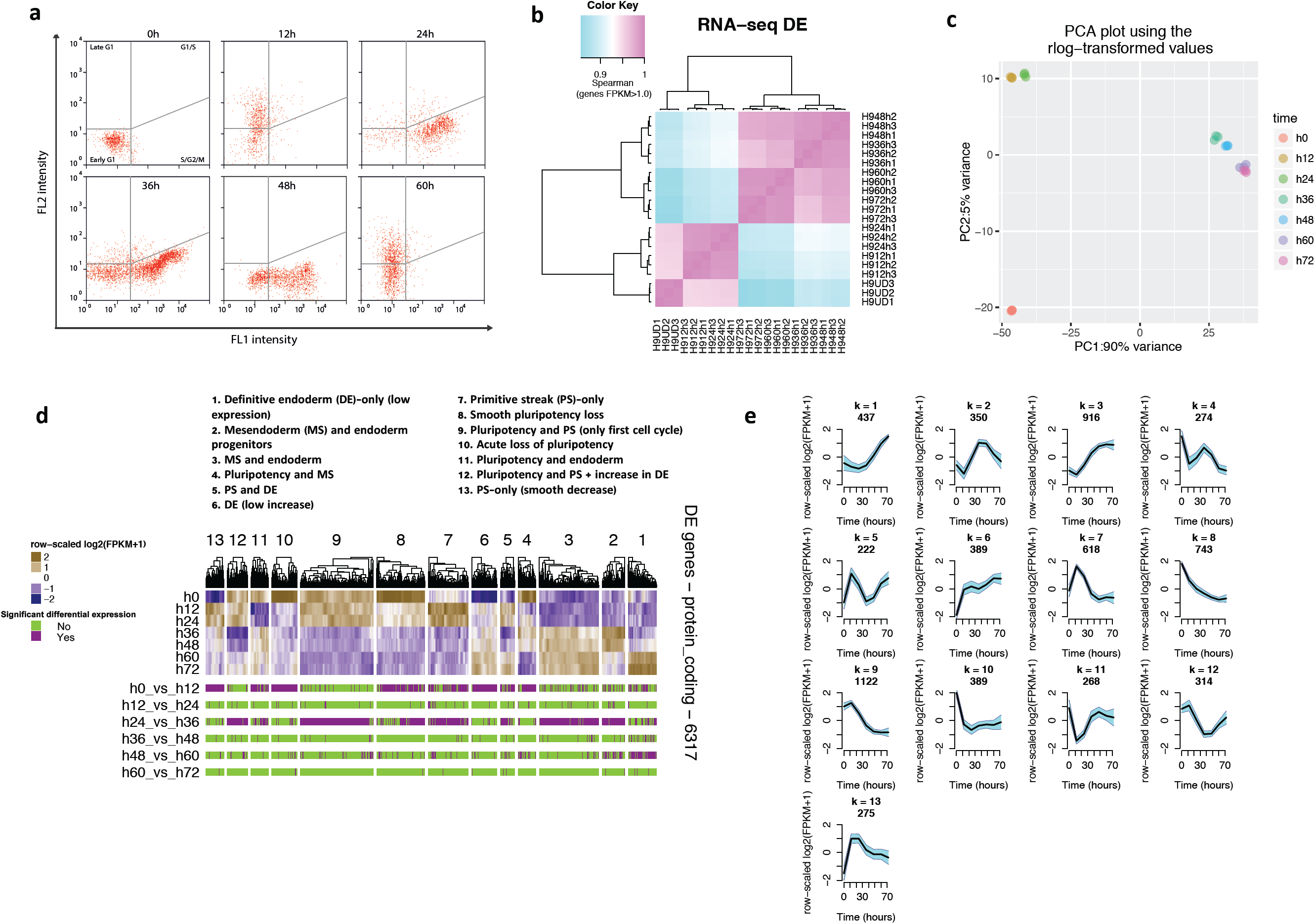
Cell cycle synchronization during differentiation of hESCs reveals novel transiently expressed early mesendoderm genes. (**a**) FACs analyses showing cell cycle progression of EG1-FUCCI hPSCs differentiating into endoderm. (**b**) Hierarchical clustering analysis pf Spearman correlation for gene expression in RNA-Seq samples. (**c**) Principal component analysis (PCA) of G1 synchronised FUCCI-hPSCs differentiating into endoderm after 12hrs (Late G1 of 1st cell cycle); 24 hr (S/G2/M of 1st cell cycle); 36 hrs (S/G2/M phase of 2nd cell cycle), 48 hrs (end of second cell cycle) and 60/72 hrs (G1 of 3rd cell cycle). (**d**) *k*-means clustering of differentially expressed genes (0h, 12h, 24h, 36h, 48h, 60h and 72h) and annotated clusters (*k*=13). Model based optimal number of clusters with 6,317 protein coding genes was computed. The number of clusters *k* was selected as the one that minimized the Bayesian Information Criterion. (**e**) Average pattern of expression of the 13 clusters identified by *k*-means clustering.

**Supplementary Figure 2:**
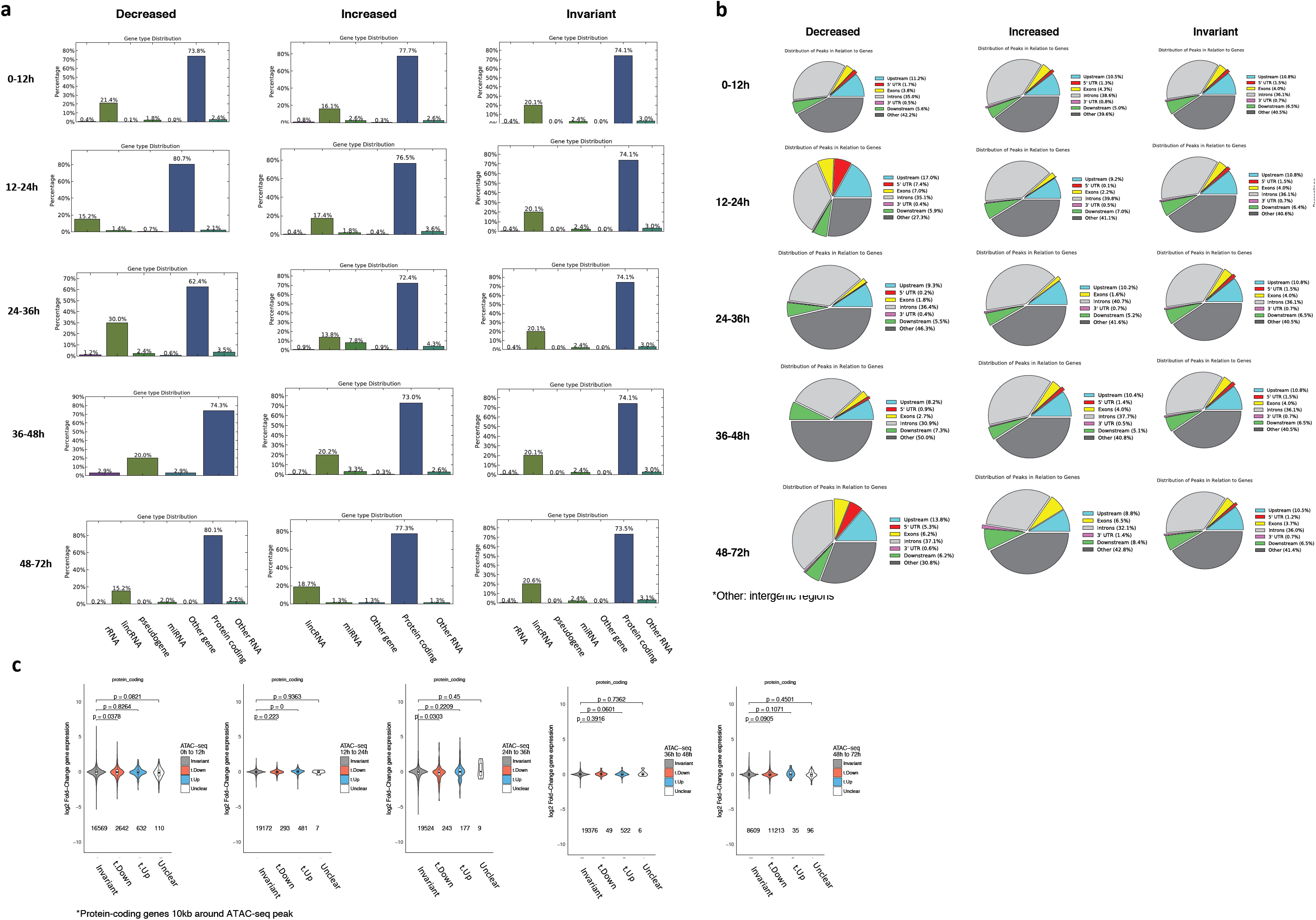
Characterization of genomic regions displaying changes in chromatin accessibility during differentiation. (**a**) Annotation of chromatin accessibility peaks to gene category during differentiation of cell synchronised EG1-Fucci hPSCs. (**b**) Annotation of chromatin accessibility peaks to intra- or inter-genic regions. (**c**) Distribution of log_2_ fold-changes in RNA expression in regions with decreased, increased, or invariant chromatin accessibility defined by ATAC-Seq. *P*-values reported by non-parametric Wilcoxon Rank Sum test. Analysis performed on protein-coding genes located 10 kb around ATAC-Seq peaks.

**Supplementary Figure 3:**
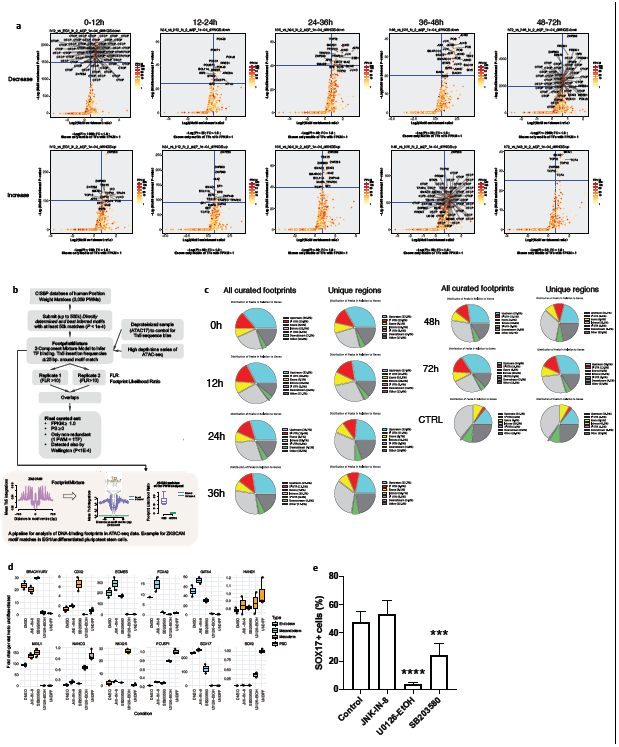
Transcription factors binding motifs in region displaying dynamic chromatin changes during endoderm differentiation. (**a**) Motif enrichment analyses in ATAC-seq peaks changing between different phases of the cell cycle. The analyses shown include only motifs for DNA-binding proteins with expression above 1 FPKM. (**b**) Computational pipeline for digital genomic footprinting in ATAC-Seq data to identify transcription factors controlling differentiation during cell cycle progression. Example for ZKSCAN1 motif matches for CisBP Position Weight Matrix M4646 at 0h. (**c**) Annotation of transcription factor footprints into intra- or inter-genic regions. A deproteinized (naked DNA) ATAC-seq sample was used a control. (**d**) Q-PCR analyses showing the effect of different inhibitors of AP1-related signalling pathways on the expression of pluripotency (NANOG, POU5F1/OCT4 and SOX2), mesoderm (BRACHYURY/T, CDX2, NKX2.5, HAND1, MIXL1) and endoderm markers (FOXA2, SOX17, GATA4, EOMES). cA1ATD hPSC line was grown for 3 days in culture conditions inducing endoderm differentiation in the presence of DMSO (control), JNK-in-8, SB 203580, U0126. cA1ATD hPSCs grown in culture condition maintaining pluripotency were used as negative control. The average of 2 independant experiments including 6 different biological replicates is provided with STD. (**e**) FACs analyses showing the expression of the endoderm marker SOX17 in cA1ATD hPSCs differentiated into endoderm in the presence of the denoted inhibitors.

## References

Adam, R.C., et al. Pioneer factors govern super-enhancer dynamics in stem cell plasticity and lineage choice. Nature. 521, 366–370 (2015)

Atlasi Y, Stunnenberg HG. The interplay of epigenetic marks during stem cell differentiation and development. Nat Rev Genet 18(11), 643–658 (2017)

Argelaguet R, Clark SJ, Mohammed H, Stapel LC, Krueger C, Kapourani CA, Imaz-Rosshandler I, Lohoff T, Xiang Y, Hanna CW, Smallwood S, Ibarra-Soria X, Buettner F, Sanguinetti G, Xie W, Krueger F, Göttgens B, Rugg-Gunn PJ, Kelsey G, Dean W, Nichols J, Stegle O, Marioni JC, Reik W. Multi-omics profiling of mouse gastrulation at single-cell resolution. Nature 576(7787), 487–491 (2019)

Buenrostro JD, Giresi PG, Zaba LC, Chang HY, Greenleaf WJ. Transposition of native chromatin for fast and sensitive epigenomic profiling of open chromatin, DNA-binding proteins and nucleosome position. Nature Methods 10, 1213–1218 (2013)

Calder A, Roth-Albin I, Bhatia S, Pilquil C, Lee JH, Bhatia M, Levadoux-Martin M, McNicol J, Russell J, Collins T, Draper JS. Lengthened G1 phase indicates differentiation status in human embryonic stem cells. Stem Cells Dev. 22(2), 279–295 (2013)

Chia CY, Madrigal P, Denil SLIJ, Martinez I, Garcia-Bernardo J, El-Khairi R, Chhatriwala M, Shepherd MH, Hattersley AT, Dunn NR, Vallier L. GATA6 Cooperates with EOMES/SMAD2/3 to Deploy the Gene Regulatory Network Governing Human Definitive Endoderm and Pancreas Formation. Stem Cell Reports 12, 57–70 (2019)

Chu LF, Leng N, Zhang J, Hou Z, Mamott D, Vereide DT, Choi J, Kendziorski C, Stewart R, Thomson JA. Single-cell RNA-seq reveals novel regulators of human embryonic stem cell differentiation to definitive endoderm. Genome Biol. 17, 173 (2016)

Corces MR, Granja JM, Shams S, Louie BH, Seoane JA, Zhou W, Silva TC, Groeneveld C, Wong CK, Cho SW, Satpathy AT, Mumbach MR, Hoadley KA, Robertson AG, Sheffield NC, Felau I, Castro MAA, Berman BP, Staudt LM, Zenklusen JC, Laird PW, Curtis C; Cancer Genome Atlas Analysis Network, Greenleaf WJ, Chang HY. The chromatin accessibility landscape of primary human cancers. Science 362, pii: eaav1898. (2018)

Cuomo ASE, Seaton DD, McCarthy DJ, Martinez I, Bonder MJ, Garcia-Bernardo J, Amatya S, Madrigal P, Isaacson A, Buettner F, Knights A, Natarajan KN; HipSci Consortium, Vallier L, Marioni JC, Chhatriwala M, Stegle O. Single-cell RNA-sequencing of differentiating iPS cells reveals dynamic genetic effects on gene expression. Nat Commun. 11(1), 810 (2020)

Dalton, S. Linking the Cell Cycle to Cell Fate Decisions. Trends Cell Biol. 25, 592–600 (2015)

De Robertis EM, Kuroda H. Dorsal-ventral patterning and neural induction in Xenopus embryos. Annu Rev Cell Dev Biol. 20, 285–308 (2004)

Dixon, J.R., et al. Chromatin architecture reorganization during stem cell differentiation. Nature. 518, 331–336 (2015)

Festuccia N, Gonzalez I, Owens N, Navarro P. Mitotic bookmarking in development and stem cells. Development 144(20), 3633–3645 (2017)

Fisher JB, Pulakanti K, Rao S, Duncan SA. GATA6 is essential for endoderm formation from human pluripotent stem cells. Biol Open 6(7), 1084–1095 (2017)

Genga RMJ, Kernfeld EM, Parsi KM, Parsons TJ, Ziller MJ, Maehr R. Single-Cell RNA-Sequencing-Based CRISPRi Screening Resolves Molecular Drivers of Early Human Endoderm Development. Cell Rep 27(3), 708–718 (2019)

González, A.J., et al. Early enhancer establishment and regulatory locus complexity shape transcriptional programs in hematopoietic differentiation. Nat Genet. 47, 1249–59 (2015)

Hnisz D, Abraham BJ, Lee TI, et al. Super-enhancers in the control of cell identity and disease. Cell 155(4), 934–947 (2013)

Li L, Wang Y, Torkelson JL, Shankar G, Pattison JM, Zhen HH, Fang F, Duren Z, Xin J, Gaddam S, Melo SP, Piekos SN, Li J, Liaw EJ, Chen L, Li R, Wernig M, Wong WH, Chang HY, Oro AE. TFAP2C- and p63-Dependent Networks Sequentially Rearrange Chromatin Landscapes to Drive Human Epidermal Lineage Commitment. Cell Stem Cell 24, 271–284 (2019)

Li QV, Dixon G, Verma N, Rosen BP, Gordillo M, Luo R, Xu C, Wang Q, Soh CL, Yang D, Crespo M, Shukla A, Xiang Q, Dündar F, Zumbo P, Witkin M, Koche R, Betel D, Chen S, Massagué J, Garippa R, Evans T, Beer MA, Huangfu D. Genome-scale screens identify JNK–JUN signaling as a barrier for pluripotency exit and endoderm differentiation. Nat Genet 51, 999–1010 (2019b)

Madrigal P, Alasoo K. AP-1 Takes Centre Stage in Enhancer Chromatin Dynamics. Trends Cell Biol 28(7), 509–511 (2018)

Malaguti M, Migueles RP, Blin G, Lin CY, Lowell S. Id1 Stabilizes Epiblast Identity by Sensing Delays in Nodal Activation and Adjusting the Timing of Differentiation. Dev Cell. 50, 462-477.e5 (2019)

Mayran A, and Drouin J. Pioneer transcription factors shape the epigenetic landscape. The Journal of Biological Chemistry 293, 13795–13804 (2018)

Na J, Furue MK, Andrews PW. Inhibition of ERK1/2 prevents neural and mesendodermal differentiation and promotes human embryonic stem cell self-renewal. Stem Cell Res 5, 157–169 (2010)

Neph S, Vierstra J, Stergachis AB, Reynolds AP, Haugen E, Vernot B, Thurman RE, John S, Sandstrom R, Johnson AK, Maurano MT, Humbert R, Rynes E, Wang H, Vong S, Lee K, Bates D, Diegel M, Roach V, Dunn D, Neri J, Schafer A, Hansen RS, Kutyavin T, Giste E, Weaver M, Canfield T, Sabo P, Zhang M, Balasundaram G, Byron R, MacCoss MJ, Akey JM, Bender MA, Groudine M, Kaul R, Stamatoyannopoulos JA. An expansive human regulatory lexicon encoded in transcription factor footprints. Nature 489, 83–90 (2012)

Oomen ME, Hansen AS, Liu Y, Darzacq X, Dekker J. CTCF sites display cell cycle-dependent dynamics in factor binding and nucleosome positioning. Genome Res 29, 236–249 (2019)

Orford K, Scadden D. Deconstructing stem cell self-renewal: genetic insights into cell-cycle regulation. Nat Rev Genet 9, 115–128 (2008)

Parker SC, Stitzel ML, Taylor DL, et al. Chromatin stretch enhancer states drive cell-specific gene regulation and harbor human disease risk variants. Proc Natl Acad Sci U S A. 110(44), 17921–17926 (2013)

Pauklin S, Vallier L. The cell-cycle state of stem cells determines cell fate propensity. Cell 155, 135–147 (2013)

Pauklin S, Madrigal P, Bertero A, Vallier L. Initiation of stem cell differentiation involves cell cycle-dependent regulation of developmental genes by Cyclin D. Genes Dev 30(4), 421–33 (2016)

Piper J, Assi SA, Cauchy P, Ladroue C, Cockerill PN, Bonifer C, Ott S. Wellington-bootstrap: differential DNase-seq footprinting identifies cell-type determining transcription factors. BMC Genomics 16, 1000 (2015)

Radzisheuskaya A, Chia Gle B, dos Santos RL, Theunissen TW, Castro LF, Nichols J, Silva JC. A defined Oct4 level governs cell state transitions of pluripotency entry and differentiation into all embryonic lineages. Nat Cell Biol. 15(6), 579–90 (2013)

Reiter F, Wienerroither S, Stark A. Combinatorial function of transcription factors and cofactors. Curr Opin Genet Dev. 43, 73–81 (2017)

Roadmap Epigenomics Consortium, et al. Integrative analysis of 111 reference human epigenomes. Nature. 518, 317–330 (2015)

Sakaue-Sawano A, Kurokawa H, Morimura T, Hanyu A, Hama H, Osawa H, Kashiwagi S, Fukami K, Miyata T, Miyoshi H, Imamura T, Ogawa M, Masai H, Miyawaki A. Visualizing spatiotemporal dynamics of multicellular cell-cycle progression. Cell 132(3), 487–498 (2008)

Séguin CA, Draper JS, Nagy A, Rossant J. Establishment of endoderm progenitors by SOX transcription factor expression in human embryonic stem cells. Cell Stem Cell 3(2), 182–95 (2008)

Slattery M, Zhou T, Yang L, Dantas Machado AC, Gordân R, Rohs R. Absence of a simple code: how transcription factors read the genome. Trends Biochem Sci. 39, 381–399 (2014)

Singh AM, Reynolds D, Cliff T, Ohtsuka S, Mattheyses AL, Sun Y, Menendez L, Kulik M, Dalton S. Signaling network crosstalk in human pluripotent cells: a Smad2/3-regulated switch that controls the balance between self-renewal and differentiation. Cell Stem Cell 10(3, 312–26 (2012)

Singh AM, Chappell J, Trost R, Lin L, Wang T, Tang J, Matlock BK, Weller KP, Wu H, Zhao S, Jin P, Dalton S. Cell-Cycle Control of Developmentally Regulated Transcription Factors Accounts for Heterogeneity in Human Pluripotent Cells. Stem Cell Reports 1(6), 532–544 (2013)

Singh AM, Sun Y, Li L, Zhang W, Wu T, Zhao S, Qin Z, Dalton S. Cell-Cycle Control of Bivalent Epigenetic Domains Regulates the Exit from Pluripotency. Stem Cell Reports 5(3), 323–336 (2015)

Singh AM, Trost R, Boward B, Dalton S. Utilizing FUCCI reporters to understand pluripotent stem cell biology. Methods 101, 4–10 (2016)

Teo AK, Arnold SJ, Trotter MW, Brown S, Ang LT, Chng Z, Robertson EJ, Dunn NR, Vallier L. Pluripotency factors regulate definitive endoderm specification through eomesodermin. Genes Dev. 25(3), 238–50 (2011)

Touboul T, Hannan NR, Corbineau S, Martinez A, Martinet C, Branchereau S, Mainot S, Strick-Marchand H, Pedersen R, Di Santo J, Weber A, Vallier L. Generation of functional hepatocytes from human embryonic stem cells under chemically defined conditions that recapitulate liver development. Hepatology 51(5), 1754–65 (2020)

Tsankov AM, Gu H, Akopian V, Ziller MJ, Donaghey J, Amit I, Gnirke A, Meissner A. Transcription factor binding dynamics during human ES cell differentiation. Nature 518, 344–349 (2015)

Vallier L, Touboul T, Brown S, Cho C, Bilican B, Alexander M, Cedervall J, Chandran S, Ahrlund-Richter L, Weber A, Pedersen RA. Signaling pathways controlling pluripotency and early cell fate decisions of human induced pluripotent stem cells. Stem Cells 27(11), 2655–66 (2009)

Wang Z, Oron E, Nelson B, Razis S, Ivanova N. Distinct lineage specification roles for NANOG, OCT4, and SOX2 in human embryonic stem cells. Cell Stem Cell 10(4), 440–54 (2012)

Wang A, Yue F, Li Y, Xie R, Harper T, Patel NA, Muth K, Palmer J, Qiu Y, Wang J, Lam DK, Raum JC, Stoffers DA, Ren B, Sander M. Epigenetic priming of enhancers predicts developmental competence of hESC-derived endodermal lineage intermediates. Cell Stem Cell. 16, 386–99 (2015)

Wang Q, Zou Y, Nowotschin S, Kim SY, Li QV, Soh CL, Su J, Zhang C, Shu W, Xi Q, Huangfu D, Hadjantonakis AK, Massagué J. The p53 Family Coordinates Wnt and Nodal Inputs in Mesendodermal Differentiation of Embryonic Stem Cells. Cell Stem Cell. 20, 70–86 (2017)

Yiangou L, Ross ADB, Goh KJ, Vallier L. Human Pluripotent Stem Cell-Derived Endoderm for Modeling Development and Clinical Applications. Cell Stem Cell 22, 485–499 (2018)

Zhang X, Cheong SM, Amado NG, et al. Notum is required for neural and head induction via Wnt deacylation, oxidation, and inactivation. Dev Cell. 32, 719–730 (2015)

Ziller, M.J., et al. Dissecting neural differentiation regulatory networks through epigenetic footprinting. Nature. 518, 355–359 (2015)

## Supplementary References

Baek S, Goldstein I, Hager GL. Bivariate Genomic Footprinting Detects Changes in Transcription Factor Activity. Cell Rep. 19, 1710–1722 (2017)

Bailey, T. et al. Practical guidelines for the comprehensive analysis of ChIP-seq data. PLoS Comput. Biol. 9, e1003326 (2013). PMID: 24244136

Bertero, A., et al. Activin/nodal signaling and NANOG orchestrate human embryonic stem cell fate decisions by controlling the H3K4me3 chromatin mark. Genes Dev. 29, 702–717 (2015).

Brons, I.G., et al. Derivation of pluripotent epiblast stem cells from mammalian embryos. Nature 448, 191–195 (2007)

Buenrostro JD et al. Transposition of native chromatin for fast and sensitive epigenomic profiling of open chromatin, DNA-binding proteins and nucleosome position. Nat Methods. 2013 Dec;10(12):1213–8.

Conesa A, et al. A survey of best practices for RNA-seq data analysis. Genome Biol. 2016;17:13. PMID: 26813401

Cremona MA, Xu H, Makova KD, Reimherr M, Chiaromonte F, Madrigal P. Functional data analysis for computational biology. Bioinformatics. 2019 Sep 1;35(17):3211-3213.PMID: 30668667

Down TA, Piipari M, Hubbard TJ. Dalliance: interactive genome viewing on the web. Bioinformatics 27:889–890 (2011) PUBMED: 21252075

ENCODE Project Consortium. An integrated encyclopedia of DNA elements in the human genome. Nature 489, 57–74 (2012)

Falcon, S. and Gentleman, R. Using GOstats to test gene lists for GO term association. Bioinformatics 23, 257–258 (2007)

Favorov, A. et al. Exploring Massive, Genome Scale Datasets with the GenometriCorr Package. PLoS Comput Biol 8, e1002529 (2012).

Feng, X., Grossman, R. & Stein, L. PeakRanger: a cloud-enabled peak caller for ChIP-seq data. BMC Bioinformatics 12, 139 (2011)

Grant CE, Bailey TL, Noble WS. FIMO: scanning for occurrences of a given motif. Bioinformatics. 2011 Apr 1;27(7):1017-8. PMID: 21330290

Gusmao EG et al. Analysis of computational footprinting methods for DNase sequencing experiments. Nat Methods. 2016 Apr;13(4):303-9. PMID: 26901649

Kim, D., et al. TopHat2: accurate alignment of transcriptomes in the presence of insertions, deletions and gene fusions. Genome Biol. 14, R36 (2013)

Ibrahim, M. M. et al. JAMM: a peak finder for joint analysis of NGS replicates. Bioinformatics 31, 48–55 (2015)

Liao, Y., et al. featureCounts: an efficient general purpose program for assigning sequence reads to genomic features. Bioinformatics 30, 923–930 (2014).

Li, H. & Durbin, R. Fast and accurate long-read alignment with Burrows-Wheeler transform. Bioinformatics 26, 589–595 (2010).

Li, Q. et al. Measuring reproducibility of high-throughput experiments. Ann. Appl. Stat. 5, 1752–1779 (2011).

Love, M., et al. Moderated estimation of fold change and dispersion for RNA-seq data with DESeq2. Genome Biol. 15, 550 (2014)

Madrigal, P. On Accounting for Sequence-Specific Bias in Genome-Wide Chromatin Accessibility Experiments: Recent Advances and Contradictions. Front. Bioeng. Biotechnol. 3, 144 (2015)

Marçais G, Kingsford C. A fast, lock-free approach for efficient parallel counting of occurrences of k-mers. Bioinformatics. 2011 Mar 15;27(6):764–70 (2011) PMID: 21217122

Mateos, J. L. et al. Combinatorial activities of SHORT VEGETATIVE PHASE and FLOWERING LOCUS C define distinct modes of flowering regulation in Arabidopsis. Genome Biol. 16, 31 (2015)

Pauklin, S. and Vallier, L. The cell-cycle state of stem cells determines cell fate propensity. Cell 155, 135–147 (2013)

Pauklin, S., et al. Initiation of stem cell differentiation involves cell cycle-dependent regulation of developmental genes by Cyclin D. Genes Dev. 30, 421–33 (2016)

Piper J, Elze MC, Cauchy P, Cockerill PN, Bonifer C, Ott S. Wellington: a novel method for the accurate identification of digital genomic footprints from DNase-seq data. Nucleic Acids Res. 2013 Nov;41(21):e201. PMID: 24071585

Piper, J., et al. Wellington-bootstrap: differential DNase-seq footprinting identifies cell-type determining transcription factors. BMC Genomics 16, 1000 (2015)

Quinlan, A. R. & Hall, I. M. BEDTools: a flexible suite of utilities for comparing genomic features. Bioinformatics 26, 841–842 (2010)

Ramsay JO, Silverman BW. Functional Data Analysis. New York: Springer; 2005.

Robinson, M.D., et al. edgeR: a Bioconductor package for differential expression analysis of digital gene expression data. Bioinformatics 26, 139–140 (2010)

Sakaue-Sawano et al., 2009 Visualizing spatiotemporal dynamics of multicellular cell-cycle progressions with fucci technology. Cell. 2008 Feb 8;132(3):487-98. PMID: 18267078

Schep, A.N. et al. Structured nucleosome fingerprints enable high-resolution mapping of chromatin architecture within regulatory regions. Genome Res. 25, 1757–1770 (2015). PMID: 26314830

Shen, L., et al. diffReps: Detecting Differential Chromatin Modification Sites from ChIP-seq Data with Biological Replicates. PLoS ONE 8, e65598 (2013)

Sung, M.H, et al. DNase footprint signatures are dictated by factor dynamics and DNA sequence. Mol Cell 56, 275–85 (2014)

Vallier, L., et al. Early cell fate decisions of human embryonic stem cells and mouse epiblast stem cells are controlled by the same signalling pathways. PLoS One 4, e6082 (2009)

Weirauch MT, et al. Determination and inference of eukaryotic transcription factor sequence specificity. Cell 158, 1431–1443 (2014)

Yardimci, G.G., et al. Explicit DNase sequence bias modeling enables high-resolution transcription factor footprint detection. Nucleic Acids Res. 42, 11865–11878 (2014).

